# Endothelial Cell Autophagy Suppresses Metastasis In Mouse Mammary and Pancreatic Neuroendocrine Tumor Models

**DOI:** 10.64898/2026.01.28.702341

**Authors:** Nancy León-Rivera, Brayden Chin, Ashley Quintana, Sofia Bustamante-Eguíguren, Anthony Gacasan, Monica Nanni, Jayanta Debnath, Teresa Monkkonen

## Abstract

Autophagy, a key lysosomal degradation pathway regulating metabolic adaptation in cancer, plays fundamental roles in both the tumor and host stromal compartments during cancer progression. An important unanswered question is whether and how autophagy in specific host stromal elements, such as endothelial cells, influences metastasis. Here, we scrutinize how the genetic loss of autophagy in endothelial cells impacts primary tumor progression and metastasis in the Polyoma Middle T (*PyMT*) model of luminal B breast cancer. In both autochthonous and orthotopic mammary transplant models, *PyMT* primary tumor growth is significantly delayed upon endothelial cell Atg12 or Atg5 genetic deletion (*Atg12 or 5* ECKO), which correlates with increased tumor cell apoptosis and HIF1α activation. In contrast, *PyMT*-bearing *Atg12* ECKO mice exhibit increased metastasis, as well as higher rates of primary tumor and lung metastatic recurrence following surgical resection of *PyMT* primary tumors. Experimental metastasis assays further corroborate that loss of endothelial cell autophagy in *Atg12* ECKO host animals promotes PyMT metastatic colonization and outgrowth, resulting in increased lung metastases compared to controls. Similarly, in the Rat Insulin Promoter T antigen pancreatic neuroendocrine tumor (RT2-PNET) model, endothelial cell deletion of *Atg12* promotes liver micro-metastases. Taken together, these results from distinct preclinical cancer models reveal that endothelial cell autophagy suppresses metastatic seeding and progression and broach that autophagy inhibition in host endothelial cells may adversely influence the efficacy of systemic autophagy-lysosomal pathway inhibition in the clinical oncology setting.

## Introduction

A hallmark of tumor progression is a dramatic change in the metabolic pathways used and metabolic needs[1]. One cellular process that promotes metabolic adaptation is macroautophagy, hereafter termed autophagy, a highly conserved lysosomal degradation process employed by cells to degrade and recycle proteins or other cellular components in response to stress or starvation. Abundant evidence supports that autophagy is frequently upregulated in cancer cells exposed to stressors such as rapid tumor growth and chemotherapy treatment[2, 3]. In many cancers, including breast cancer, this increase in autophagic flux supports the advanced stages of tumor progression[2]. As a result, several autophagy inhibitors remain under active investigation, such as antimalarials that disrupt lysosomal acidification, and newer agents that target different proteins required for autophagy, including palmitoyl protein thioesterase 1 (PPT1), and the Unc51-like (ULK1/2 kinases)[2, 4]. Indeed, numerous clinical trials have examined whether autophagy inhibition is beneficial for the treatment of breast and other cancers and a recent clinical trial employed hydroxychloroquine as a potential agent to reduce disseminated tumor cell burden in breast cancer patients with the goal of preventing metastatic recurrence[5].

While many pre-clinical studies focus on the role of tumor cell autophagy[2, 3, 6], autophagy in host stromal cells has also emerged as a potential important regulator of the tumor microenvironment[7–9]. Early studies showed that non-tumor cell autophagy supported RAS-driven tumorigenesis in a *Drosophila* model[10]. Follow-up work corroborated that the loss of autophagy in host fibroblasts or stellate cells limited the primary tumor growth in diverse solid tumor models[8]. Non-tumor cell autophagy in host stromal cells may contribute to successful tumor metabolic adaptation by supplying key amino acids to tumor cells[9]. In addition, the autophagy machinery may control non-degradative functions in the tumor microenvironment and other contexts by facilitating the secretion of cytokines[7, 8, 11], type I collagen[8], and extracellular vesicles[12]. Finally, many studies collectively suggest the importance of autophagy in regulating signaling mechanisms governing immune cell recruitment[13].

Autophagy has been previously studied in endothelial cells, in both non-cancer and cancerous contexts[14, 15]. Mice with conditional knockout of *Atgs* required for autophagosome formation demonstrate that autophagy mediates fundamental vascular homeostasis processes such as removal of cholesterol[16], cellular alignment in response to shear stress[17], activation of nitric oxide signaling[18], and packaging and release of Von Willibrand Factor with injury[19, 20]. GFP-LC3 autophagy reporter activity has been localized to sites of neutrophil transendothelial migration in a model of acute ischemia, and can regulate CD31 via non-canonical LC3-associated phagocytosis[21]. Mammalian target of rapamycin (mTOR), an upstream regulator of autophagy, has also been associated with cytoprotective effects in endothelial cells during ischemia/ hypoxia conditions[22, 23]. In tumors, the associated vasculature plays important roles in tumor physiology, regulating nutrient access, hypoxia, necrosis, drug delivery, and immune infiltration[1, 24]. Accordingly, previous studies have examined the loss of endothelial cell autophagy in the B16F10 model of melanoma. Upon conditional endothelial cell knockout of *Atg5*, B16F10 melanomas exhibited increased vascular tortuosity and density, while pericyte coverage, vessel size, and perfusion are all attenuated[25]. These angiogenic phenotypes are associated with reduced primary tumor growth, but no changes in metastasis[25]. Other studies reveal distinct impacts of chloroquine on tumor angiogenesis and angiogenic signaling versus genetic depletion of autophagy[25, 26]. For example, in HUVECs, chloroquine treatment alters the transcription and signaling associated with ANGP1, PDGF, and EDN1, which is not observed upon *Atg5* knockdown, consistent with an autophagy-independent for chloroquine in the control of vascular function[25, 26]. Loss of tumor endothelial cell autophagy is also correlated with increased CD8+ T cell infiltration into melanoma, potentially via upregulation of STING and increased endothelial cell expression of the ICAM and VCAM adhesion molecules[27]. Treatment with the MTOR inhibitor, everolimus (which stimulates autophagy), reduces immature and mature tumor vasculature *in vivo* without altering VEGF levels or vascular permeability[28], while combined MTOR and VEGF inhibition in pancreatic neuroendocrine tumors disrupts metabolic symbiosis between hypoxic and normoxic tumor cells and slows tumor growth[29]. While data from these in vivo melanoma models support important roles for endothelial cell autophagy in primary tumor growth, the broader role of endothelial cell autophagy in other cancers, including its specific effects on metastasis, largely remains uncertain.

Here, we scrutinize the genetic loss of endothelial cell (EC) autophagy in the progression and metastasis of PyMT-driven luminal B breast cancer and pancreatic neuroendocrine tumors (PNET). Similar to previous work in melanoma, the systemic genetic ablation of *Atg5/12* in endothelial cells in mice slows primary tumorigenesis in both autochthonous and orthotopic transplantation models of Polyoma Middle T (*PyMT*) mammary tumorigenesis. On the other hand, genetic loss of *Atg12* in endothelial cells results in enhanced *PyMT* metastasis to lungs, increased primary tumor relapse and lung metastasis following the resection of primary *PyMT* tumors, and increased levels of *PyMT* lung metastatic colonization following tail-vein injection. Moreover, while endothelial cell loss of *Atg12* in the Rat Insulin Promoter T Antigen pancreatic neuroendocrine tumor (*RT2-PNET*) model has minimal effects on primary PNET growth or progression; increased spontaneous micro-metastases to the liver is observed. In contrast to the melanoma transplantation models previously employed in studies of EC autophagy, results in these two distinct autochthonous cancer models broach a previously unrecognized functional role for EC autophagy in suppressing metastasis.

## Methods

### Mouse models and maintenance

All mice were maintained and experiments performed according to UCSF IACUC standards under animal protocol AN170608 or SDSU IACUC standards under animal protocol 25-084.

### Atg 12 and Atg5 ECKO mice

Compound strains for endothelial specific autophagy deletion were produced by intercrossing Cdh5CreER[30] mouse (B6.FVB-Tg(Cdh5-cre)7Mlia/J) with conditional CRE recombinase activity under the VE cadherin promoter to those with conditional floxed alleles for ATG5^F/F^; Atg5^tm1Myok^ (MGI# 5784717)[31] (obtained from N. Mizushima) or ATG12^F/F^; Atg12^tm1.1Jdth^ (MGI# 5784706)[32]. These mice also carried a ubiquitously expressed RFP reporter allele Gt(ROSA)26Sor^tm1Hjf^ (MGI# 3696099) with a Rosa26 promoter driven lox-stop-lox cassette preceding RFP[33]. *Atg12 fl/fl; Rfp/Rfp* only female mice were interbred to *Cdh5CreER; Atg12 fl/fl; Rfp/Rfp* males to generate *Atg12 ECKO* (with a similar scheme to generate Atg5 ECKO). Genotyping was performed using published primers and protocols[34], or Transnetyx automated genotyping services. To initiate recombination, mice were administered tamoxifen (Sigma) at 0.0002 mg/kg dissolved in peanut oil (Sigma) by oral gavage for 5 consecutive days as previously described[34]. Peanut oil was used as a vehicle control.

### RT2-PNET2 *mice*

Previously generated *RT2-PNET2* mice[35] (MGI# 2429941) were bred to the *CdhCreER; Atg12 fl/fl; Rfp/Rfp* mice. The normal mouse diet was supplemented with optional sucrose rich pellets in caging (Envigo catalog no.TD8649) starting at 10 weeks of age[35]. Tumor volumes were calculated using 0.5* width^2^ * length and measured by caliper. To initiate recombination, mice were administered tamoxifen (Sigma) at 0.0002 mg/kg dissolved in peanut oil (Sigma) by oral gavage for 5 consecutive days as previously described[34]. Peanut oil was used as a vehicle control. Tamoxifen or peanut oil were administered starting at 5-6 or 10 weeks of age.

### PyMT organoid collection and orthotopic transplantation

Female *MMTV-PyMT*; Tg(MMTV-PyVT)634Mul/LellJ mice (Jackson Labs Stock# 022974)[36] in the C57/B6 background were originally obtained from Z. Werb (UCSF). Organoids were collected from endpoint animals when the largest tumor has volume 2000 mm^3^ using 0.5 * width^2^ * length formula. Tumors were monitored two times per week until reaching 1000 mm^3^ volume by caliper, and subsequently monitored daily until reaching volume of 2000 mm^3^ (or other humane endpoints). Mice were harvested and organoids prepared as previously[37] and stored in liquid nitrogen.

Prior to injection, cells are plated at 0.5-1 vial per plate and cultured in growth media consisting of DMEM/F12 (Gibco) with 5% FBS (Atlas), 50 ng/mL EGF (Peprotech), 1 mg/mL hydrocortisone (Sigma), 1% Pen/Streptomycin (ThermoFisher), and 10 mg/mL insulin (Sigma cat no. I1882) for 3 days prior to collection before digestion in 0.025% Trypsin (Gibco). Cells are counted with a Countess, and injected in 50% Matrigel (BC Biosciences, Lot 15899) and 50% dPBS (Gibco). All cells were cultured in 5% CO_2_ in a humidified incubator at 37°C. One week prior to tumor cell transplantation, female transplant recipient mice 8-12 weeks in age were treated for 5 consecutive days with tamoxifen or peanut oil (vehicle) as detailed above to allow for *Atg12* recombination in host endothelial cells; thereafter transplantation/injection of PyMT organoids into the mammary gland was performed according to Deome et al[38]. All lungs were harvested under isofluorane.

### PyMT resection study

*PyMT* organoids were transplanted as above ∼one week after the last oral gavage as detailed above. Tumors were monitored and resected at a size of 1000 mm^3^ by electrocautery under anesthesia, and then animals harvested after a ‘chase’ period of 50 days post-resection unless animals met other endpoint criteria prior to the completion of the chase period.

### Autochthonous PyMT ECKO

*Cdh5CreER; Atg12 fl/fl; Rfp/Rfp; PyMT+* males were crossed to *Atg12 fl/fl; Rfp/Rfp* females to generate *PyMT+* female progeny. Mice were treated with tamoxifen or peanut oil as above with the first day of oral gavage being the first palpable detection of a tumor. Animals were harvested when the largest tumor reached an endpoint of 2000 mm^3^.

### Experimental metastasis assays

*PyMT* primary organoids were cultured ∼3 days in defined media (see above) prior to preparation for lateral tail vein injection. One week prior to tail vein injection, syngeneic female recipient mice 6 to 12 weeks of age were treated with either tamoxifen or peanut oil (vehicle) by oral gavage daily for 5 days at doses above to achieve recombination in endothelial cells. Thereafter, 5 x10^5^ live PyMT cells/ 150 μL dPBS were injected by 25 gauge needle into syngeneic female recipient mice. Mice were harvested under isofluorane 1 month post injection.

### Flow cytometry

Murine lungs were minced and digested in 10 mL of PBS with 2.5 U/mL Dispase, 1 mg/mL Collagenase II (Gibco), and 15 U/mL DNAse (Stem Cell) at 37 degrees for 30’ −1 hour shaking at 200 RPM for each animal. 5 mL of Endothelial cell growth media (EGM) (Lonza) were added, the cells strained through a 100uM filter, and red blood cell lysis was performed for 1’ using 1 mL lysis buffer (Sigma) before adding 15-20 mL EGM, and cells were spun down at 400 RCF for 5 minutes. Cells were resuspended and incubated in Fc blocker (Miltenyi), 1:10 in 5% FBS/ dPBS in polystyrene tubes for 10’ on ice prior to incubation with CD31-FITC antibody (Biolegend Clone 390 #102406), at 1:50 on ice for 30’. Antibody was washed out, and Sytox blue (Invitrogen) used as a live/ dead marker at concentration 1:500. Analysis was performed on a BD LSRFortessa X-20. At least 10,000 singlet live cells were analyzed per animal. Gating and analysis performed using FloJo software.

### Histology

Tissues were fixed in 3.7% aqueous zinc buffered paraformaldyhyde (Z-fix, Anatech) at 4 degrees C on a shaker for ∼24 hours, then moved to 70% EtOH at 4 degrees C on a shaker for >24 hours, then processed and embedded. Sections were cut at a thickness of 4-5 μm. All hematoxylin and eosin staining was performed by the UCSF Mouse Pathology Core. For quantification of lung metastasis, 3 step sections with >50 μm distance between were taken and stained with hematoxylin and eosin. Stitched H&E images of 40x were produced by whole slide scanning with an Aperio XT Whole Slide Scanner (Leica), and areas of lungs and metastases defined manually using QuPath software. Necrosis was quantified in 1-2 hematoxylin and eosin stained tumor cross sections in QuPath manually, by including largely eosinophilic areas characterized by cellular and nuclear debris.

### Immunohistochemistry

Slides were dewaxed in a series of xylenes, ethanols of decreasing concentration, and washed in PBS. Antigen retrieval was performed in a Biocare Medical Antigen Retrieval Chamber decloaker in DAKO Target Retrieval Solution or Bio SB Immuno DNA Retriever 20X with Citrate, and slides cooled for 20 minutes. Blocking buffer consisted of 10% goat/ horse serum in PBS plus 0.1% Tween20 (Sigma) (hereafter, PBST). Primary antibody staining was done in blocking buffer either overnight at 4 degrees C, or 1 hour at room temperature on a rocker. Please see supplemental table 1 for primary antibody company and concentration information. Washes were performed in PBST. Secondary antibody staining was done using a biotinylated secondary antibody (Vector Biotinylated Anti mouse/rabbit IgG, reconstituted at 1 mg/mL) diluted 1:500 in PBS for 1 hour. After washing, the Vector Labs ABC Elite HRP kit was used, followed by Vector Nova Red Substrate Kit for 5-20’. Counterstaining was done for ∼30 seconds in Hematoxylin QS (Vector Labs). Quantification of immunohistochemistry images were performed using ImageJ software (version 1.8). TIFF images of stained CK14 were calibrated to pixels and for each image the total tissue area was first defined manually using color threshold function in HSB (Hue, Saturation, Brightness), which was used to determine total stained tumor area in pixels. The same thresholding was then applied to isolate positively stained CK14 areas and quantified in pixels and expressed as a percentage of the total tissue area. To validate reproducibility, a subset of images were re-analyzed to confirm accuracy of results.

### Immunofluorescence

Similar protocols were followed for IHC, with secondary antibodies replaced by 1:500 AlexaFluor secondary antibodies diluted in PBS. Nuclear counterstain was done by 2 ng/mL Hoechst (3343, Thermo Scientific) in PBS for 12 minutes at room temperature. After washes, slides were mounted in Vectashield Mounting Media for Fluorescence. See Supplementary Table 1 for further details on primary antibody concentrations, company and catalog numbers. Ten blinded images per tumor acquired with Olympus BX63 with CellSens Standard software, were selected primarily from regions of viable, non-necrotic tissue. Quantification of IF staining was performed using QuPath software. Cytokeratin 5 (CK5) mean fluorescence intensity (MFI) was quantified using Image J (version 1.8). For each tumor, 10 random representative images with minimal necrosis were taken at identical exposures, converted to 8-bit greyscale, and the CK5 positive channel was quantified as above for CK14. To examine vessel size, we used CD31 immunofluorescence images and ImageJ, together with an adapted macro as described previously[39].

### Blood glucose levels

BGL was obtained at 14 weeks of age using a Contour Next EZ Blood Glucose with corresponding test strips, and obtained at a similar time of day (2:00 pm) to minimize circadian effects.

### RT2-PNET micrometastasis

Livers of mice from the early deletion at 5 weeks of age (see above) were obtained for FFPE (see above), sectioned, and stained by IHC. One section of >2 lobes of liver was quantified for SV40+ cells identified by immunohistochemistry using QuPath software. Area of livers was measured using Q path to normalize number of cells.

### Statistics

Statistical analysis was performed using the statistical test in figure legends or results text. Survival curves generated in GraphPad Prism, and analyzed by log rank test. Otherwise, data analyzed via GraphPad Prism by ANOVA analysis with Tukey’s test for multiple comparisons or student’s t test, with Mann Whitney or Welch’s correction as noted. We eliminated outliers for the resection lung metastasis study and the autochthonous ECKO study using the ROUT test for outliers with Graphpad Prism.

## Results

### Genetic ablation of ATG12 in endothelial cells slows primary tumor growth but enhances lung metastasis in an autochthonous mammary cancer model

Previous studies demonstrate that the conditional loss of endothelial cell (EC) autophagy via conditional endothelial knockout of *Atg5* alters angiogenesis in B16F10 melanoma tumors in mice[25]. Tumors derived from B16F10 melanoma cells transplanted into host animals lacking EC autophagy display increased vascular tortuosity and density, decreased vessel size and perfusion, and decreased pericyte coverage, together with reduced tumor volume[25]. To determine how endothelial autophagy influences tumorigenesis in an autochthonous murine tumor model, mice with floxed conditional ablation alleles targeting autophagy related protein 12 (ATG12)[32] were interbred with mice carrying the inducible *Cdh5CreER* vascular endothelial cadherin (Vascular Endothelial Cadherin) promoter driven expression of Cre fused to a mutant estrogen receptor[30] to generate *Atg12* ECKO animals. These mice also possess a ubiquitously expressed Cre-dependent red fluorescent protein (RFP) reporter[33]. We verified that *Atg12* ECKO mice displayed evidence of appropriate endothelial cell Cre-dependent knockout. We performed FACS analysis to determine the percentage of CD31 positive lung endothelial cells displaying RFP positivity. One week following vehicle or tamoxifen treatment to induce Cre activation, 75.5% of CD31+ cells from *Atg12* ECKO mice displayed RFP positivity, while *Atg12 fl/fl* controls with vehicle (mean, 0.6%), *Atg12 fl/fl* control with tamoxifen (1.0%), and *Cdh5CreER; Atg12 fl/fl* controls with vehicle (17%) all displayed significantly fewer RFP+ CD31+ cells (p ≤ 0.0001 all groups versus *Atg12* ECKO, ANOVA with Tukey’s test) (Fig.1a). See Supplemental Fig. 1 for FACS gating strategy. α-P62/SQSTM1 immunofluorescence revealed increased accumulation in CD31+ cells in *Atg12* ECKO endothelium relative to controls (Fig. 1b).

**Figure 1.**
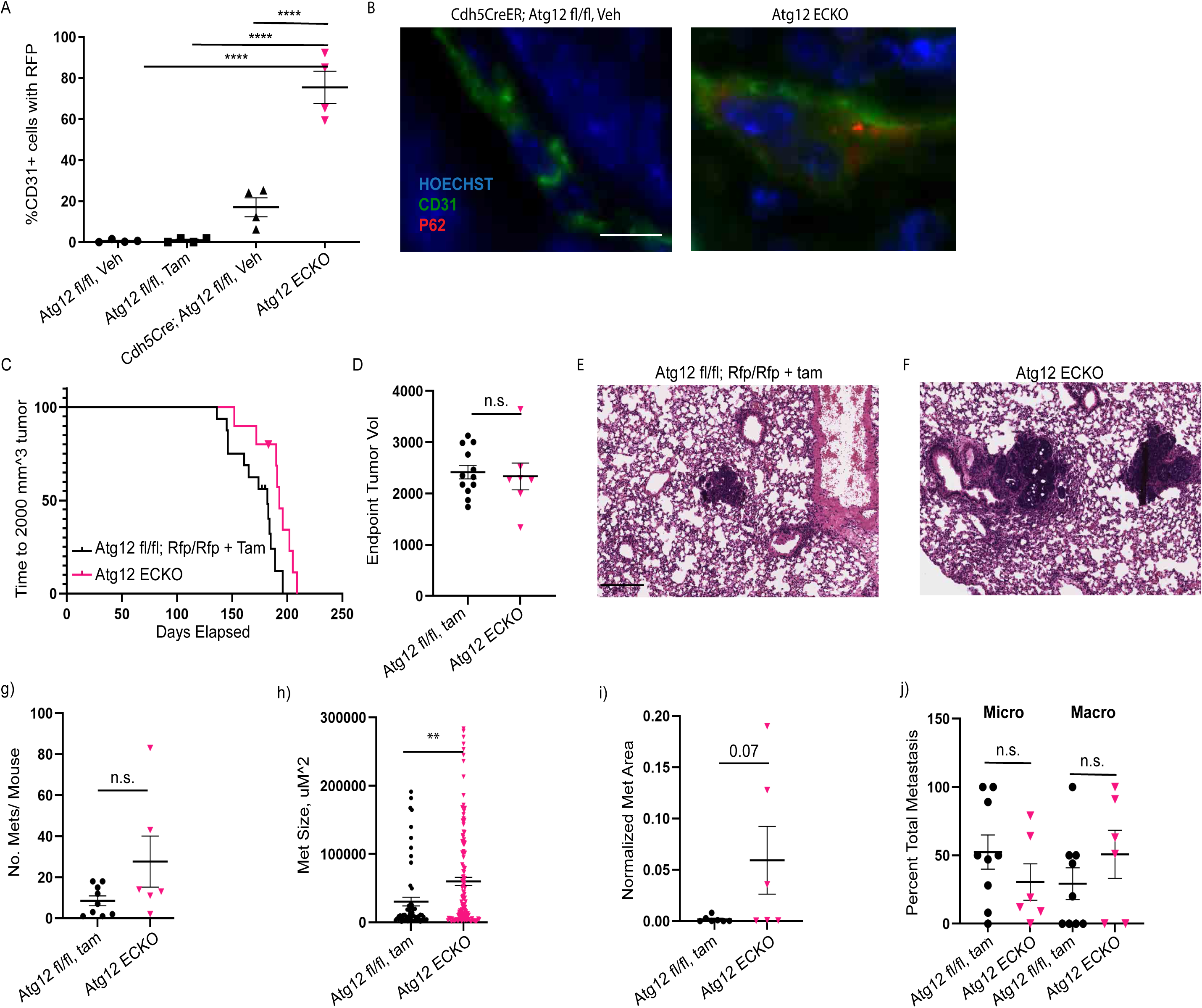
Endothelial cell autophagy KO slows autochthonous *PyMT* growth and increases lung metastasis size. (**A**) FACS sorted lung endothelial cells of adult mice show enhanced Cre-dependent RFP expression in CD31+ lung endothelial cells versus Cre- or peanut oil treated animals. One data point per mouse. (**B**) Representative P62 and CD31 stained endothelial cells showing accumulation in *Atg12* ECKO. Scale bar = 10 μM. (**C**) Autochthonous tumor growth showing statistically significant slower time to endpoint for *Atg12* ECKO mice. Controls: n= 6 *Atg12 fl/fl* mice treated with peanut oil; n= 8 *Atg12 fl/fl* mice treated with tamoxifen, and n=2 *Cdh5CreER; Atg12 fl/fl* mice treated with peanut oil. *Atg12* ECKO n= 10. (**D**) Volume of harvested *Atg12* ECKO tumors is similar. (**E-F**) Representative H and E of lung metastases from *Atg12 fl/fl* mice treated with tamoxifen and *Atg12* ECKO lung metastases. Scale bar = 200 uM (**G**) Number of lung metastases/ mouse is not different in ECKO mutants. (**H**) Sizes of all lung metastases from end stage mice. (**I**) Total metastatic area normalized to lung area per mouse. (**J**) No difference in % micrometastases (≤10,000 uM^2^) or macrometastases (>50,000 uM^2^) between groups. All quantification except f): 1 data point= 1 mouse; f) one data point= one tumor. ** indicates p≤0.01. **** indicates p≤ 0.0001.

First, we tested whether the slowed primary tumor growth phenotype observed upon loss of EC autophagy in B16F10 melanomas was corroborated in an autochthonous model of tumorigenesis^10^. Hence, we leveraged the commonly used *PyMT* model of luminal B breast cancer which exhibits high similarity to human disease[40, 41]. Importantly for these studies, we employed the PyMT tumor model generated in the C57Bl/6 genetic background, which we and other previously have shown to significantly slow the latency to tumor formation and reduce the extremely aggressive growth and progression kinetics characteristic of the FVB/n genetic background commonly utilized in studies of PyMT tumorigenesis[8, 34]. As a result, by employing a slower progression model compared to B16F10 melanomas, we hypothesized that we would be better poised to assess potential impacts of EC autophagy on both primary tumor growth and spontaneous metastasis in these models.

*Atg12 fl/fl; Rfp/Rfp* females were crossed to *Cdh5CreER; Atg12 fl/fl; Rfp/Rfp; MMTV-PyMT* males and oral gavage with tamoxifen or vehicle (peanut oil) was administered beginning the first day a palpable tumor was detected. Notably, *Cdh5CreER; Atg12 fl/fl; Rfp/Rfp* (*Atg12* ECKO) mice treated with tamoxifen displayed a longer time to endpoint tumor volume (largest tumor measuring 2000 mm^3^) versus littermate control *Atg12 fl/fl; Rfp/Rfp* mice treated with tamoxifen (p ≤ 0.0096, log rank test) (Fig. 1c). Because total tumor burden at harvest showed no significant difference in *Atg12* ECKO versus controls (p ≤ 0.71, Mann Whitney test) (Fig. 1d), we were able to scrutinize lung metastasis between these cohorts. We quantified lung metastasis in this model from H&E-stained step sections of all lobes of the lungs. All control and *Atg12* ECKO mice displayed lung metastasis. Representative H&E images show increased metastasis size in *Atg12* ECKO compared to tamoxifen treated controls (Fig. 1e, f). Although the total number of lung metastases was not significantly different between groups (Fig. 1g) (p ≤ 0.2338, Mann Whitney test), the size of individual lung metastases were significantly increased in *Atg12* ECKO mice (Fig. 1h) (p ≤ 0.0022, Mann Whitney test), broaching a potential role for EC autophagy in promoting colonization and outgrowth at the metastatic site. Additionally, when normalized to lung area, the total metastatic area in *Atg12* ECKO mice compared to controls showed a trend toward increased metastatic burden (Fig. 1i) (p ≤ 0.07, Mann Whitney test). To further scrutinize the degree of metastatic outgrowth, we categorized all lesions as either micro (≤10,000 uM^2^) or macrometastases (>50,000 uM^2^) as previously[34], but found no significant difference in the proportion of metastases between these groups (Fig. 1j) (micro: p ≤ 0.3417; macro: p ≤ 0.1319, Mann Whitney test).

### Endothelial Cell Autophagy Restricts Local and Metastatic Recurrence Following Primary Polyoma Middle T (PyMT) Tumor Excision

We next tested whether these differences in primary tumor progression and metastasis observed in autochthonous *Atg12* ECKO polyoma middle T (PyMT) tumors were recapitulated in spontaneous metastasis assays that allow for synchronous assessment of primary tumor growth across cohorts following orthotopic transplantation of tumor cells at defined timepoints. For these studies, we employed an orthotopic transplantation model using *MMTV-PyMT* derived organoids, in which five hundred thousand *PyMT* organoids were injected into *Atg12* ECKO or *Atg12 fl/fl* female mice one week after treatment with tamoxifen to allow for endothelial-specific *Atg12* deletion. As before, time to primary tumor endpoint (2000 mm^3^) volume was significantly increased in *Atg12* ECKO versus *Atg12 fl/fl* controls treated with tamoxifen (log rank test, p ≤ 0.01930) (Fig. 2a). Median survival increased from 56 (*Atg12 fl/fl* with tamoxifen) to 65 (*Atg12* ECKO) days. Growth curves of individual tumors over time show that the slowest growing tumors are *Atg12* ECKO mice (Fig. 2b). End stage H&E images showed overall similar histology in end stage *Atg12* ECKO tumors versus controls (Fig. 2c).

**Figure 2.**
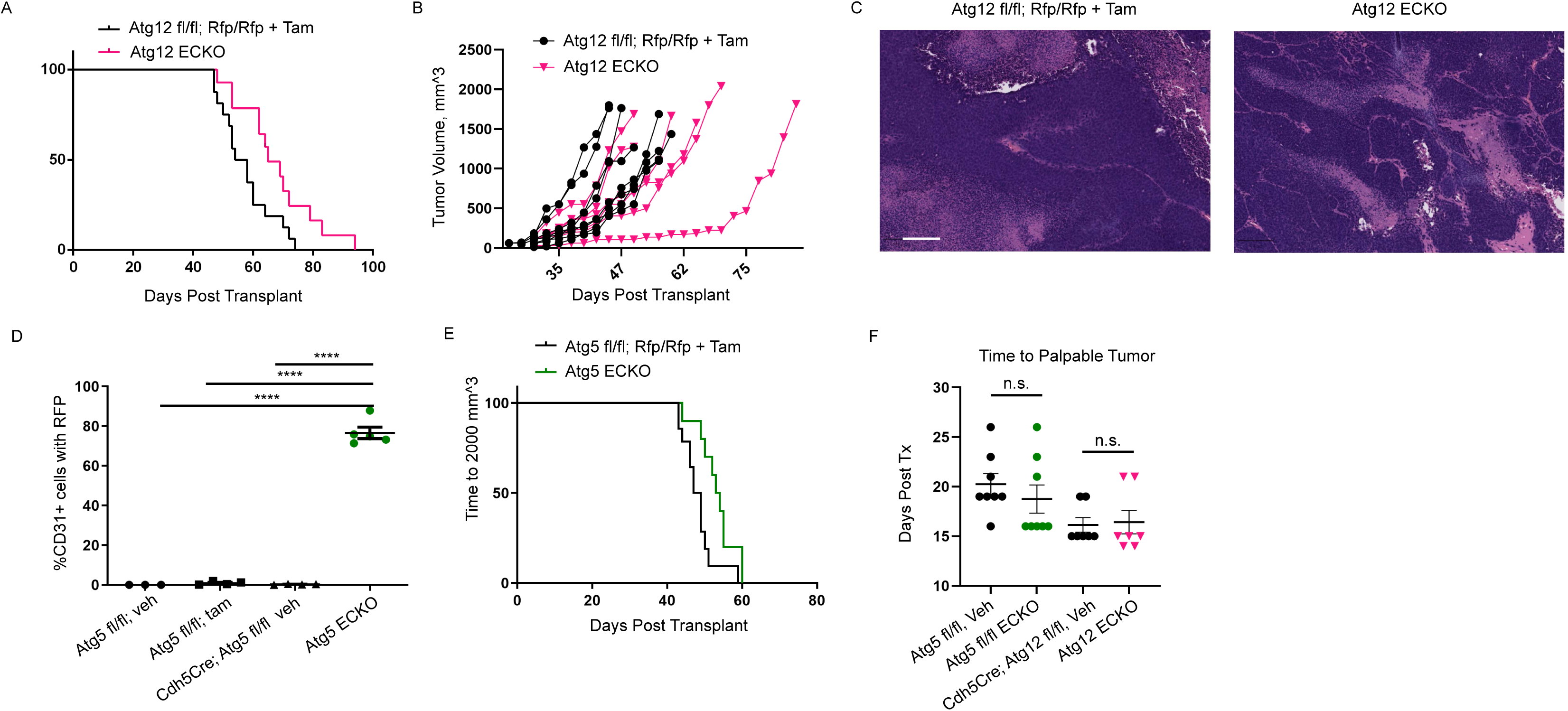
Loss of endothelial cell autophagy slows *PyMT* mammary tumor growth in vivo. (**A**)*Atg12* ECKO animals with orthotopically transplanted *PyMT* cells display longer time to tumor endpoint (2000 mm^3^) versus *Atg12 fl/fl* mice treated with tamoxifen. (**B**) Tumor volumes of individual control and *Atg12* ECKO tumors over time. Scale bar =100 μM. (**C**) Representative H and E images from control and *Atg12* ECKO end stage tumors. (**D**) FACS sorted lung endothelial cells of adult mice show enhanced Cre-dependent RFP expression in CD31+ lung endothelial cells versus Atg5 fl/fl; Rfp/Rfp or peanut oil vehicle treated animals. One data point per mouse. (**E**)*Atg5* ECKO mice display significantly longer time to 2000 mm^3^ versus control mice. (**F**) Time to detection of a palpable tumor is unaltered in *Atg5* ECKO mice or *Atg12* ECKO mice versus controls (Mann Whitney analysis). **** indicates p≤0.0001.

To confirm that our findings observed with PyMT tumor cell orthotopic transplantation into hosts lacking ATG12 in endothelial cells were independently observed upon deletion of an independent ATG in host endothelium, we generated *Atg5* ECKO mice by intercrossing *Atg5 fl/fl* mice[42] with strains carrying the inducible *Cdh5CreER* and the Cre-dependent RFP reporter. Using the experimental strategy for *Atg12* ECKO described above, we verified the *Atg5* ECKO model (Fig. 2d). *Atg*5 ECKO CD31^+^ lung cells were 76.7% RFP+, while other control groups displayed significantly fewer RFP+ cells (with vehicle: 0.02%, *Atg5 fl/fl* with tamoxifen: 0.9%, and *Cdh5CreER; Atg5 fl/fl* with vehicle: 0.3%; all controls versus *Atg*5 ECKO p ≤ 0.0001 by ANOVA with Tukey’s test). Moreover, immunohistochemistry corroborated increased levels of P62/SQSTM1 in *Atg5* ECKO endothelial cells (Supplemental Fig. 2).

Thereafter, we injected *PyMT* cells into *Atg5* ECKO mice and *Atg5 fl/fl* controls treated with tamoxifen and observed very similar results to *Atg12* ECKO, including significant increase in time to tumor endpoint (p ≤ 0.0165, log rank test) and an increase in median survival from 47 to 53 days in *Atg5* ECKO mice (Fig. 2e). To test whether tumor initiation was altered, time to detection of a palpable tumor was compared in the *Atg12* and *Atg5* ECKO studies above. *Atg12* ECKO mice had similar time to the detection of a palpable tumor versus controls (p ≤ 0.69, Mann Whitney test), as did *Atg5* ECKO mice (p ≤ 0.3239, Mann-Whitney test) (Fig. 2f). Based on these results obtain upon genetic deletion of two independent ATGs, we construe that the reduced primary tumor growth observed in these orthotopic transplantation assays arise from a general defect in EC autophagy.

To better understand the impact of EC autophagy on spontaneous metastasis, we next performed resection experiments controlling for primary tumor burden by surgically excising primary tumors from the various cohorts at matched sizes and earlier in progression (1,000 mm^3^). Eight- to twelve-week-old *Cdh5CreER; Atg12 fl/fl; Rfp/Rfp* mutants and *Atg12 fl/fl; Rfp/Rfp* littermate control mice (lacking Cre) were treated with tamoxifen for 5 consecutive days to induce recombination in Cre expressing animals. One week later, *MMTV-PyMT*-derived organoids from late stage *MMTV-PyMT* tumors (2000mm^3^) were orthotopically transplanted to ECKO mutant and control mice. Similar to our previous studies of tumor cell autophagy[34], primary mammary tumors were resected at a fixed tumor volume of ∼1000mm^3^, followed by a 50 day ‘chase’ period (schematic, Fig. 3a). At this timepoint, *Atg12* ECKO mice showed a higher frequency of both primary tumor recurrence (5/14, 36%), an increased incidence of lung metastasis (8/14, 53%), as well as increased lung metastasis in the absence of primary recurrence (6/8, 75%) in comparison to Cre-controls (*Atg12 fl/fl; Rfp/Rfp* + tamoxifen) (5/18, 28% recurrence; 6/18, 33% lung metastasis, and 4/13, 31% lung metastasis, no recurrence) (Fig. 3b). We also asked whether the time to reach 1000mm^3^ at the time of tumor resection differed between these groups, but observed no significant differences between *Atg12* ECKO and control mice (p≤0.3259, student’s t-test with Mann Whitney) (Fig. 3c). After the 50 day period, *Atg12* ECKO mice with recurred tumors had comparably sized tumors relative to *Atg12 fl/fl; Rfp/Rfp* + tamoxifen controls (p ≤ 0.9048, student’s t-test with Mann-Whitney) (Fig. 3d). Quantification of lung metastasis showed high variability. We scrutinized lung metastasis that arose in animals only in the absence of primary recurrence. There was a significant increase in the number of lung metastases per mouse in *Atg12* ECKO mice relative to *Atg12 fl/fl; Rfp/Rfp* + tamoxifen controls (p ≤ 0.0480, student’s t test/ Mann Whitney) (Fig. 3e). Furthermore, compared to controls, *Atg12* ECKO mice exhibited significantly larger average metastatic size per mouse (p ≤ 0.0437, student’s t test/Mann Whitney) (Fig. 3f) and a significant increase in normalized total metastatic area (Fig. 3g) (p ≤ 0.0237, Student’s t-test/Mann-Whitney). We found no difference in the percentage of total metastasis of micro or macrometastasis between these cohorts (Fig. 3h) (micro: p ≤ 0.4667; macro: p ≤ 0.7524, student’s t test/ Mann Whitney).

**Figure 3.**
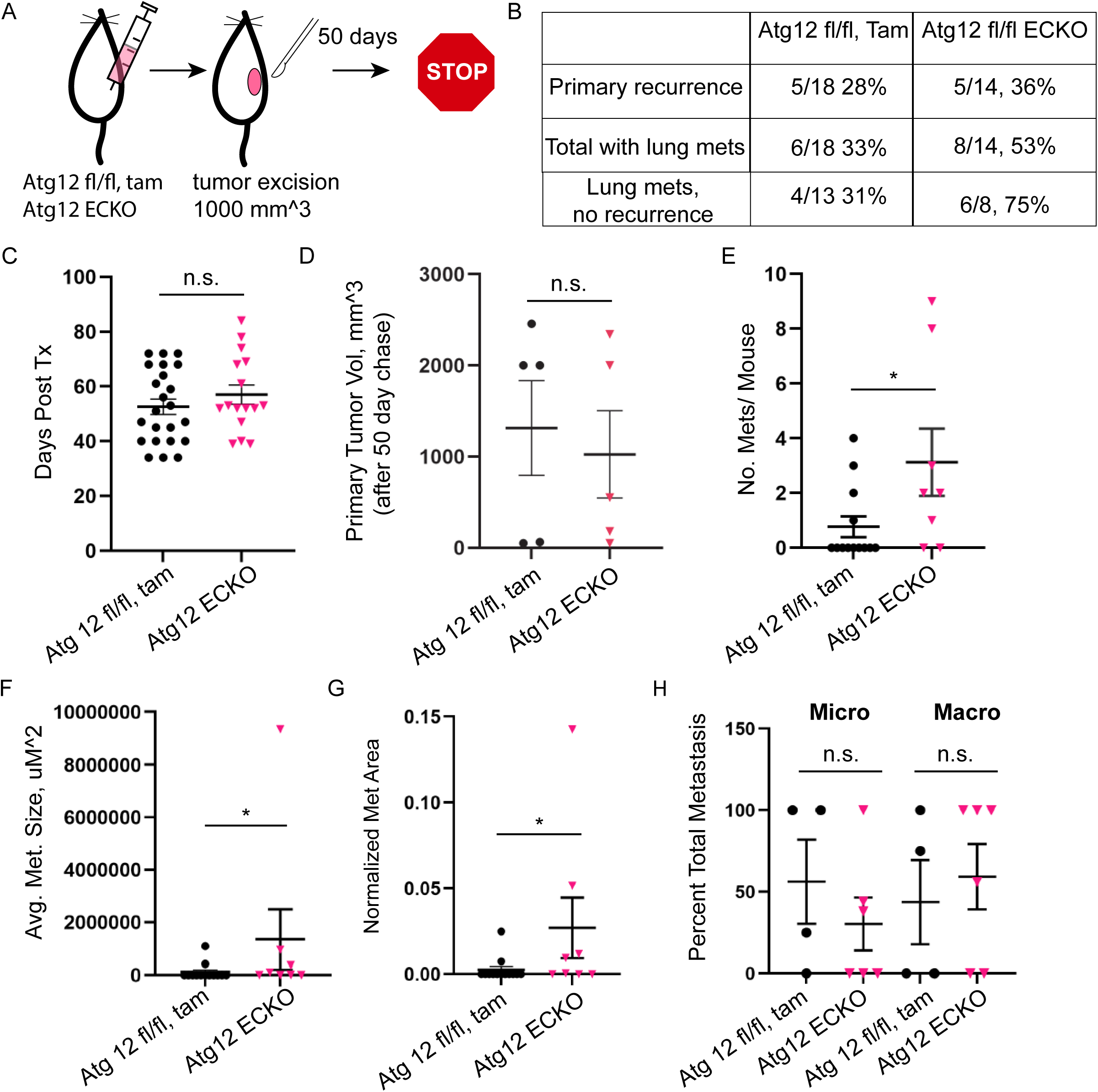
*Atg12* ECKO increases primary tumor recurrence and lung metastasis after *PyMT* resection. (**A**) Schematic of resection experiment. (**B**) *Atg12* ECKO mice display increased metastasis to lungs versus *Atg12 fl/fl* mice treated with tamoxifen, and higher rates of metastasis to lungs in the absence of primary tumor recurrence. *Atg12* ECKO mice also display a modest increase in primary tumor recurrence relative to *Atg12 fl/fl* mice treated with tamoxifen. (**C**) Time to 1000 mm^3^ is not different in *Atg12* ECKO mice. (**D**) Recurrent primary tumor volume is unaltered in *Atg12* ECKO mice versus controls with recurrence (non-recurring volumes excluded). (**E**) Number of lung metastases per mouse is significantly increased in *Atg12* ECKO mice without primary tumor recurrence. (**F**) Average metastasis size/ mouse is significantly larger in *Atg12* ECKO mice versus controls without primary tumor recurrence. (**G**) Total metastatic area/ mouse normalized to lung area is increased in *Atg12* ECKO mice without primary tumor recurrence (**H**) No significant difference in percent micro (<10,000 μM^2^) or macro lung metastases in ECKO lungs (>50,000 μM^2^). * indicates p≤0.05.

### Endothelial Cells Loss of ATG12 Promotes HIF1α Activation In PyMT Tumor Cells

To determine whether EC autophagy inhibition influences tumor cell proliferation or death, we performed immunohistochemistry and immunofluorescence staining for phospho-histone H3 (PHH3), cleaved caspase-3 (CC3), and Ki67 on tumors from *Atg12* ECKO and *Atg12 fl/fl* + tamoxifen control mice. At endpoint (∼2000mm^3^), no significant difference in the number of PHH3+ cells was found between groups (p ≤ 0.0947, student’s t test/ Mann Whitney) (Fig. 4a). Similarly, CC3 staining revealed no significant difference in tumor apoptosis in *Atg12* ECKO mice (unpaired t test with Welch’s correction, p ≤ 0.2841) (Fig. 4b). Similarly, quantification of necrotic tumor areas in end stage tumor cross sections showed no significant difference between *Atg12* ECKO and *Atg12 fl/fl* + tamoxifen controls (Fig. 4c). We also examined the resected tumors (∼1000mm^3^) to analyze an earlier stage of tumor progression. While KI67 immunofluorescence staining showed no change in proliferation compared to controls (p ≤ 0.1812, student’s t test/ Mann Whitney, outliers excluded by ROUT test) (Fig. 4e), there was a significant increase in apoptotic CC3+ cells in *Atg12* ECKO tumors versus controls (p ≤ 0.0147, student’s t-test/ Mann-Whitney) (Fig. 4f). We surmise that this increase in apoptosis during the intermediate stages of tumor development may contribute to slower primary tumor progression and longer time to tumor endpoint in *Atg12* ECKO mice (Fig.1-2).

**Figure 4.**
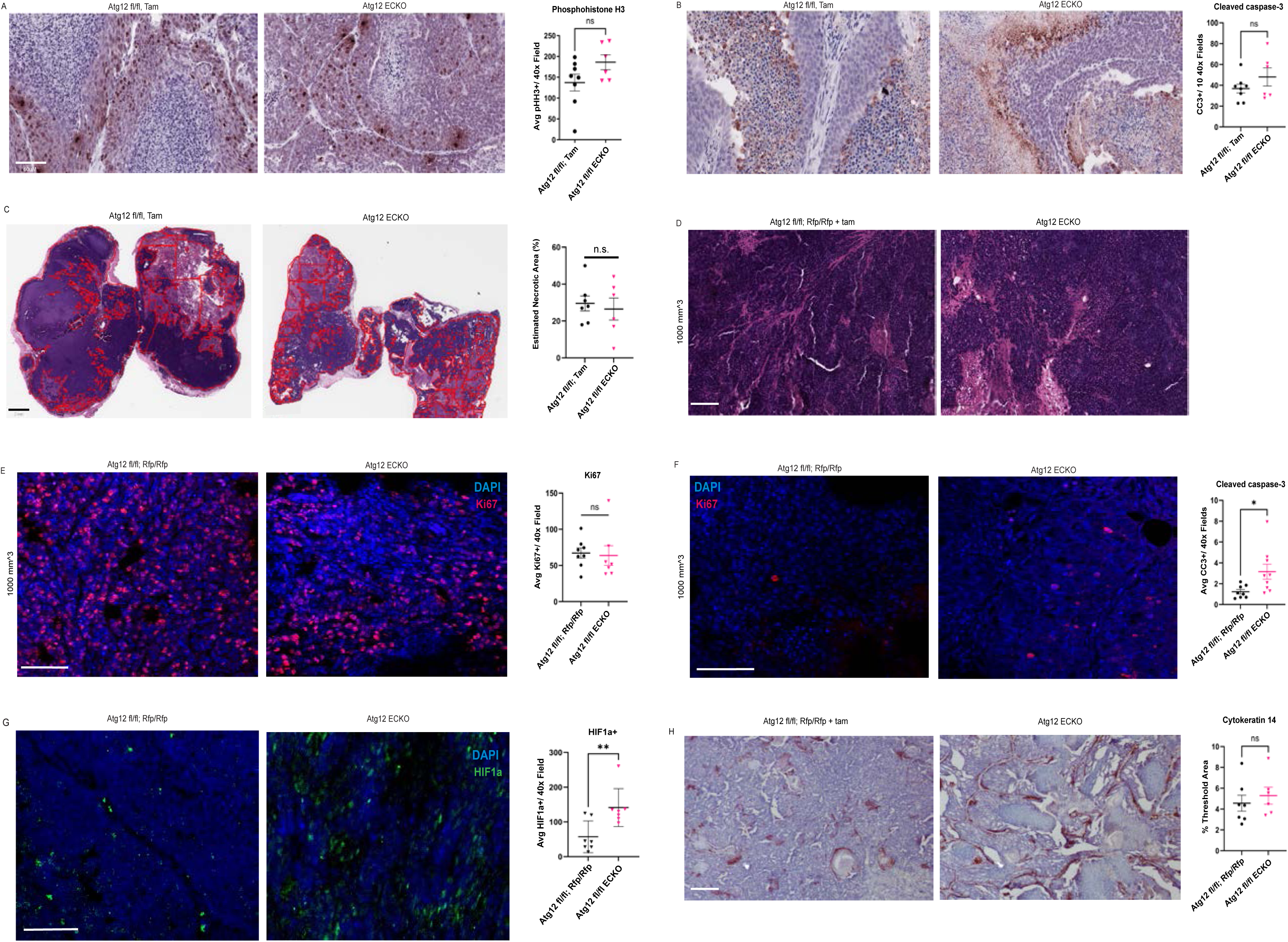
*Atg12* ECKO mice display transiently increased apoptosis, and a late-stage increase in hypoxia. Representative images and quantification of (**A**) PHH3+ cells/ 40x field in endpoint *PyMT* tumors (2000mm^3^) of *Atg12 fl/fl* + tamoxifen control and *Atg12* ECKO mice. (**B**) CC3+ cells/ 40x field in endpoint tumors. (**C**) No difference in necrotic area in tumor cross sections. Scale bar = 2 mm. (**D**) Representative H and E images of *Atg12* ECKO and control *PyMT* tumors at resection (1000mm^3^). Scale bar = 100 μM. (**E**) KI67+ cells/ 40x field in ECKO and control tumors at 1000mm^3^. (**F**) % CC3+ cells in ECKO and control tumors at 1000mm^3^ (**G**) Increased nuclear HIF1-alpha cells/ 40X field is present in endpoint ECKO *PyMT* tumors. (**H**) Images and thresholded CK14 signal/ 20X field in endpoint *PyMT* tumors is not different. One data point = one mouse. (**E-H**): scale bar = 50 μM. * indicates p ≤ 0.05; ** p ≤ 0.01.

Notably, defects in endothelial function and tumor angiogenesis can activate HIF1A (Hypoxia-Inducible Factor 1-alpha), a master regulator of the hypoxic transcriptional response, which is associated with increased metastasis to distant sites by promoting the adaptation of tumor cells to low-oxygen environments and driving pro-metastatic lineage transitions[43–45]. Accordingly, we quantified nuclear HIF1α in tumor cells, which revealed that *Atg12* ECKO tumors had higher nuclear HIF1α compared to controls (p ≤ 0.0091, student’s t test/ Mann Whitney) (Fig. 4g), consistent with an increased hypoxia-driven transcriptional response in tumors lacking EC autophagy. Of note, although increased metastasis due to loss of tumor cell autophagy has been observed previously[34], the genetic loss of tumor cell autophagy did not impact nuclear HIF1a in end stage orthotopically injected *PyMT* tumors lacking ATG 12 (Supp. Fig 3). Because aggressive subpopulations of cells exhibiting basal cytokeratin expression are associated with increased metastasis in breast cancer models, including those arising in the setting of autophagy deficiency [46], we assessed whether endothelial autophagy loss expanded CK14/CK5 positive populations in *PyMT* tumors. However, no differences in cytokeratin 14 (CK14) or expression were observed between *Atg12* ECKO and *Atg12 fl/fl* with tamoxifen controls (p ≤ 0.3660, student’s t test/ Mann Whitney) (Fig. 4h). Similarly, there were no differences in cytokeratin 5 (CK5) expression in end stage *Atg12* ECKO tumors versus controls (p ≤ 0.4848, student’s t test/ Mann Whitney) (Supplemental Fig. 4).

To determine if vascular changes associated with *Atg12* ECKO may drive the tumor phenotypes, we next examined tumor vasculature properties. End stage *Atg12* ECKO tumors revealed a significant increase in CD31+ cell density relative to controls (p ≤ 0.0004, student’s t test/ Mann Whitney) (Fig. 5a). In contrast, individual vessel size (p ≤ 0.2343) was not different (student’s t test/ Mann Whitney) (Fig.5b). We also tested whether endothelial cell proliferation or apoptosis were impacted in end stage tumors, and found increased PHH3 positivity in CD31+ cells of end stage PyMT tumors (Fig. 5c) (p ≤ 0.0022, student’s t text/ Mann Whitney). While most cell types show decreased proliferation with loss of autophagy given its homeostatic functions[8], and since endothelial cells in B16F10 melanomas displayed no change in proliferation[47], we hypothesize this increase is a response to the HIF1α activation displayed by tumor cells (Fig. 4). We also tested whether apoptosis was altered in CD31+ endothelial cells, and observed very low levels of apoptosis and no difference between groups (Fig. 5d) (p ≤ 0.7316, student’s t test/ Mann Whitney).

**Figure 5.**
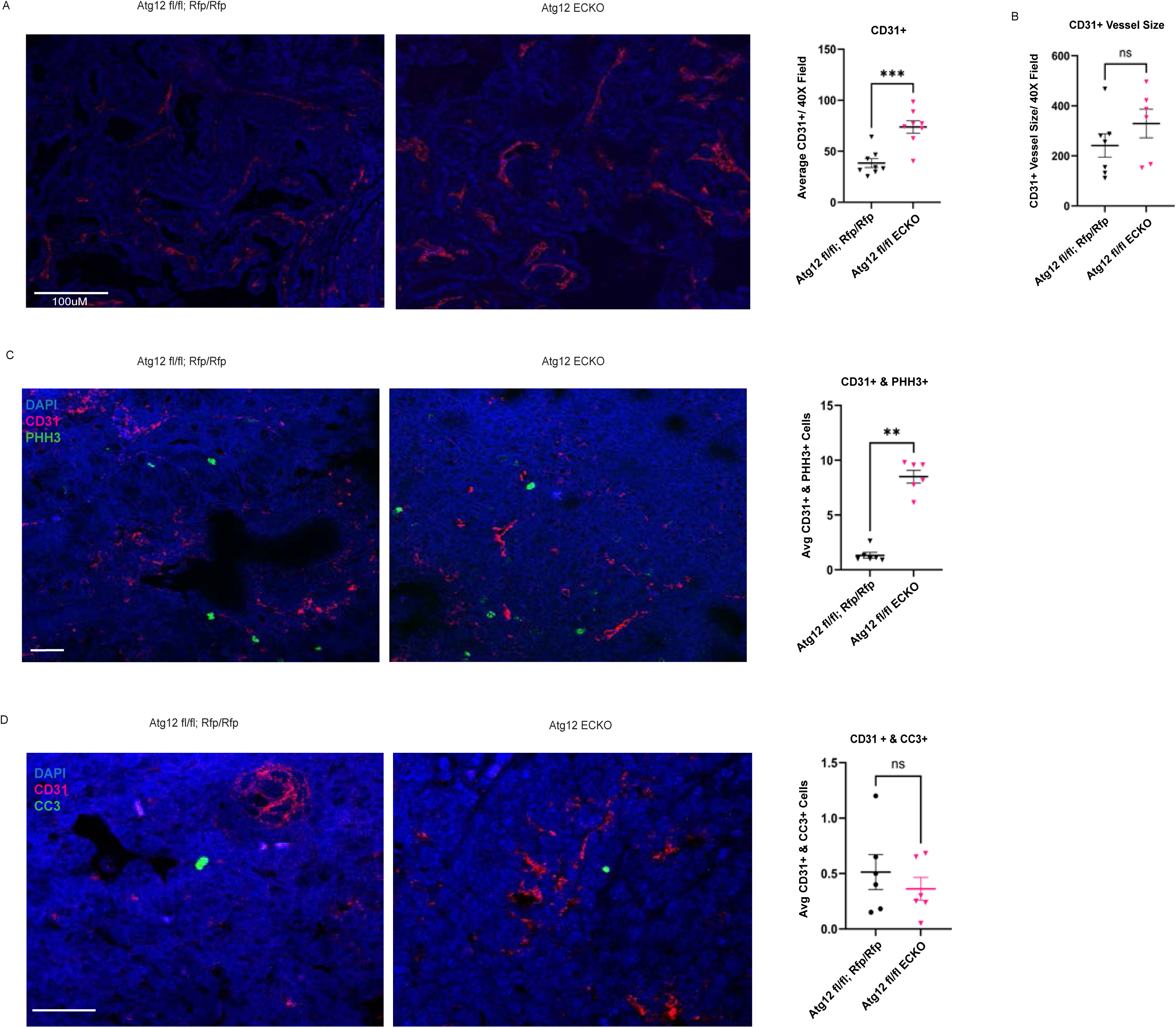
*Atg12* ECKO increases CD31+ cell density, but not vascular size/ area. (**A**) Density of CD31+ cells is increased in end stage *Atg12* ECKO tumors versus controls. Scale bar =100μM. (**B**) CD31+ vessel size in end stage *Atg12* ECKO tumors is not altered. (**C**) PHH3 immunofluorescence is increased in CD31+ cells of end stage *Atg12* ECKO tumors versus controls. (**D**) CC3 immunofluorescence is unaltered in CD31+ cells of end stage *Atg12* ECKO tumors versus controls. (**C-D**): scale bar = 50 μM ** indicates p ≤ 0.01, *** indicates p ≤ 0.001.

### Endothelial cell autophagy suppresses experimental lung metastasis

The increase in metastatic capacity we observed in *Atg12 ECKO* in both autochthonous and orthotopic tumor models motivated us to delineate whether the loss of EC autophagy impacts the later stages of the metastatic cascade, namely metastatic colonization and outgrowth, in experimental metastasis models. *PyMT*-derived tumor cells were inoculated into the lateral tail veins of vehicle-treated *Cdh5CreER; Atg12 fl/fl* mice or *Atg12* ECKO mice. As shown in representative whole lung images, *Atg12* ECKO mice exhibited an increase in the size of metastatic lesions compared to vehicle-treated, autophagy-competent controls (Fig. 6a, b). H&E staining showed no gross histological differences in representative metastases from *Atg12* ECKO mice versus controls (Fig. 6c, d). Quantification revealed an increase in the number of metastases in *Atg12* ECKO mice compared to controls (p ≤ 0.07) (Fig. 6e) as well as in the size of lung metastasis in *Atg12* ECKO versus controls (p ≤ 0.057) (Fig. 6f), although these trends were not statistically significant. Upon comparison of individual metastases across all animals in each cohort, *Atg12* ECKO mice exhibited significantly larger metastasis relative to controls (p ≤ 0.0001) (Fig. 6g). Classification of micro- and macro-metastasis revealed a significant increase in the proportion of macro-metastasis in *Atg12* ECKO mice (p ≤ 0.0431), with no significant differences in the percentage of micro-metastasis (p ≤ 0.3013, student’s t-test/Mann-Whitney) (Fig. 7h).

**Figure 6.**
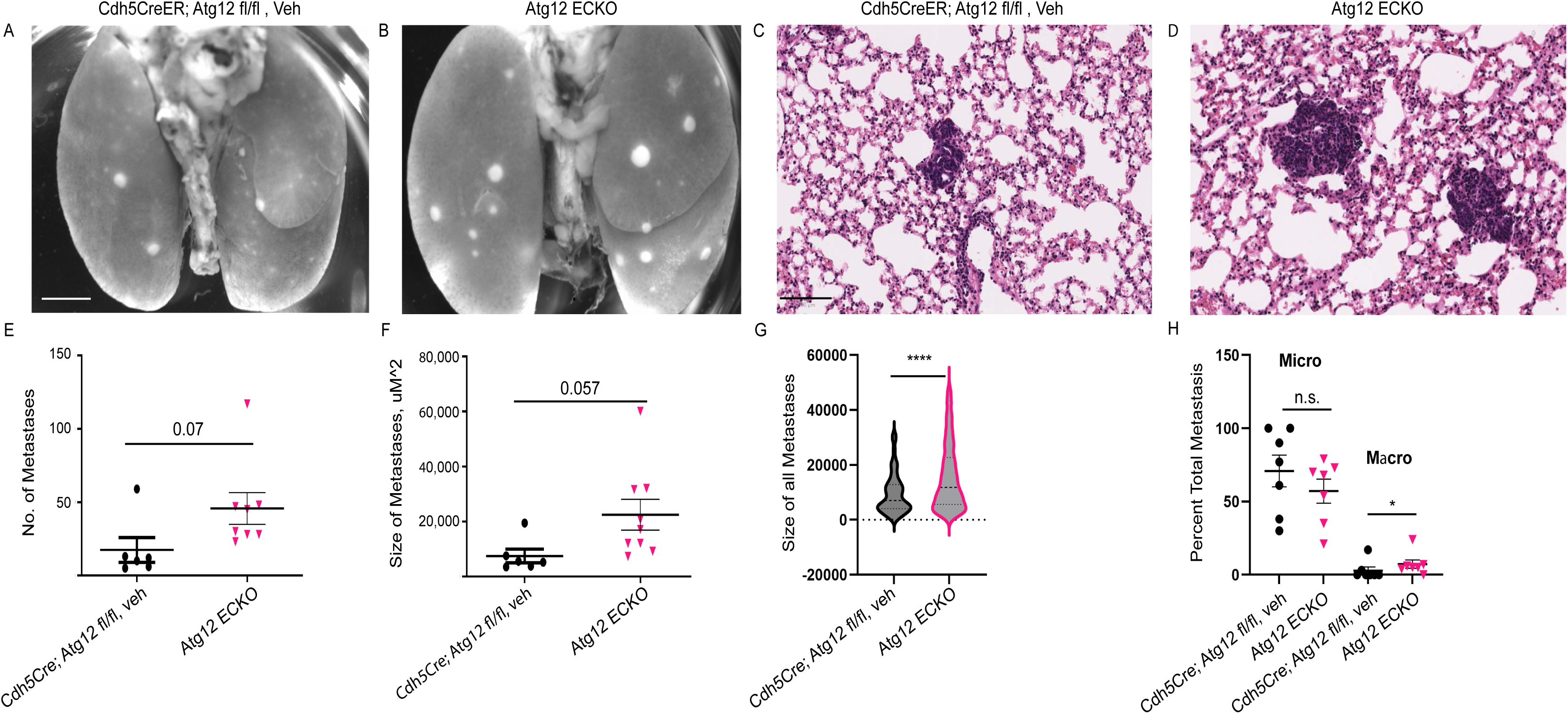
Tail Vein injected *PyMT* cells shows increased lung metastasis size in *Atg12* ECKO mice. (**A**) Representative whole lung images of *Cdh5CreER; Atg12 fl/fl* mice treated with vehicle (peanut oil) and (**B**) *Atg12* ECKO mice. Scale bar = 2 mM. Representative images of H and E stained lung metastasis from (**C**) *Cdh5CreER; Atg12 fl/fl* mice treated with vehicle and (**D**) *Atg12* ECKO lungs. Scale bar = 50 uM. (**E**) Number of metastasis may be increased in *Atg12* ECKO mice, p≤0.07. (**F**) Size of metastasis may be increased in *Atg12* ECKO mice. (**G**) Examining individual tumors (one data point= one metastasis) shows a significant increase in size of metastasis in *Atg12* ECKO mice. (**H**) Percent macrometastases is higher in *Atg12* ECKO mice relative to controls *Cdh5CreER; Atg12 fl/fl* mice treated with vehicle, while percent micrometastasis is unchanged. (D, E, G) one data point = one mouse. * indicates p ≤ 0.05, **** p≤0.0001.

**Figure 7.**
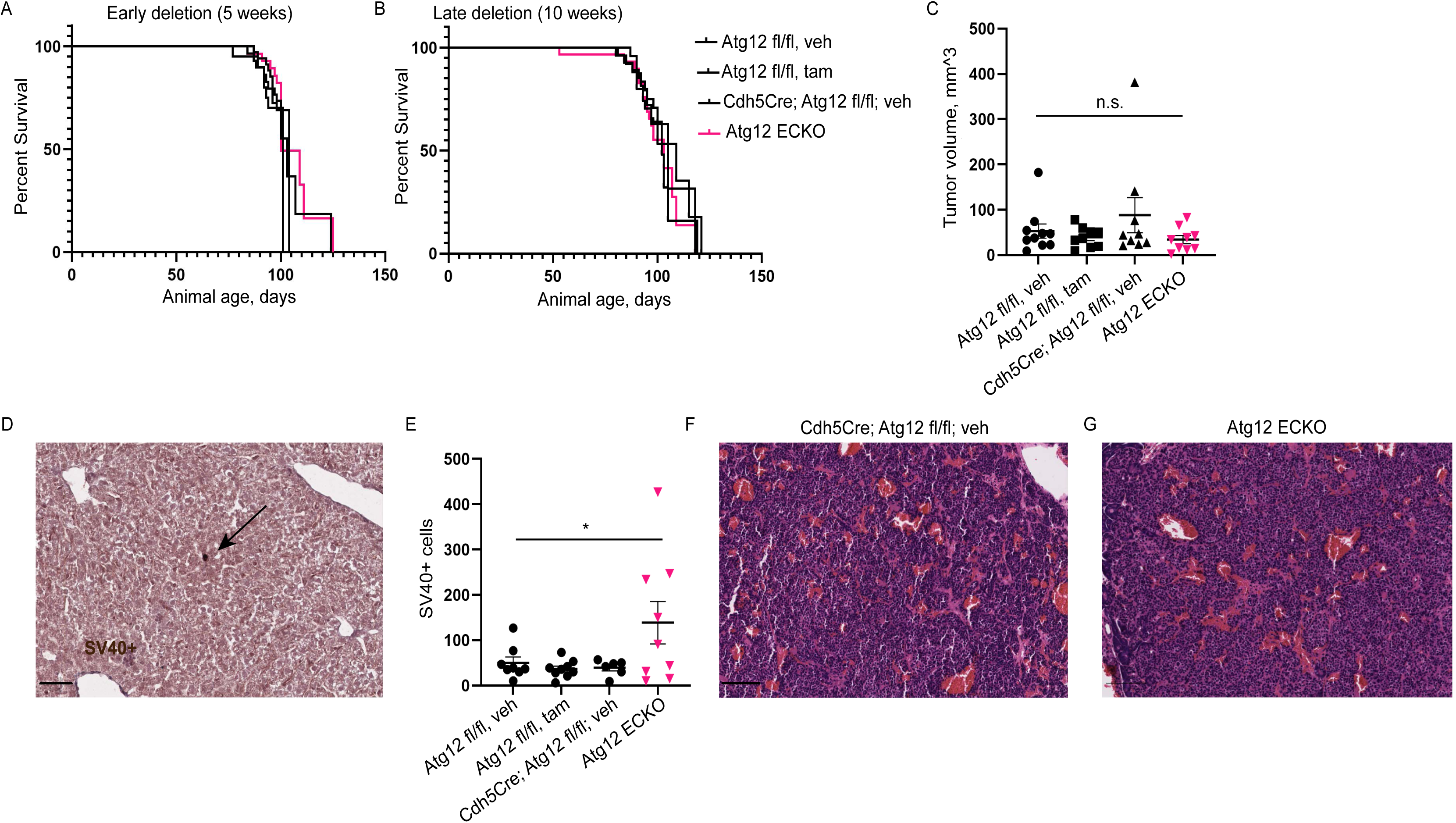
*Atg12* ECKO mice with *RT2-PNET* tumors have unaltered primary tumorigenesis and survival, with increased liver micrometastasis. (**A**) Early (5 week) EC deletion of *Atg12* does not alter survival of ECKO *RIPTAG* bearing mice. (**B**) Late *Atg12* ECKO (tamoxifen beginning at 10 weeks) does not alter *RT2-PNET* survival. A,B) log rank test (**C**) No difference in tumor volume at 14 weeks with early *Atg12* ECKO. (**D**) Example SV40+ cell in the liver. (**E**) *Atg12* ECKO livers show increased SV40+ cells versus controls. (**C,D**) one data point = one animal. Representative H and E image of (**F**) endpoint control and (**G**) endpoint *Atg12* ECKO *RT2-PNET* tumors. Scale bar = 50 uM. * indicates p ≤0.05.

### Endothelial Cell ATG12 Deletion Promotes Pancreatic Neuroendocrine Tumor Micro-metastasis to the Liver

To determine whether the observed tumor progression and metastatic phenotypes extended to a different cancer model, we examined a distinct autochthonous cancer model, the Rat insulin promoter T antigen mouse model of pancreatic neuroendocrine tumors (*RT2-PNET*), which is considers a gold-standard transgenic model for the study of tumor angiogenesis[35]. In this model, SV40 Large T antigen oncogene is expressed in pancreatic insulin-producing beta cells, resulting in a well-characterized and stereotypic time course of tumor progression, in which mice develop hyperplasic islets, followed by angiogenic islets, and ultimately, islet carcinomas at 10-12 weeks of age[35]. These mice exhibit a well-characterized angiogenic switch prior to the onset of islet carcinoma, with mice succumbing to disease at ∼14 weeks of age. Notably, because previous work shows that RT2-PNETs are more vascular and respond better to anti-angiogenic therapy relative to *PyMT* tumors[48], we interrogated whether EC autophagy functionally impacted progression prior to the crucial angiogenic switch in this model. However, early deletion of *Atg12* at 5-6 weeks of age did not alter survival in *RT2-PNET* mice with *Atg12* ECKO (Log-rank test, p ≤ 0.4276) (Fig. 7a). To confirm the presence of Cre+ endothelial cells at tumor endpoint, we performed RFP immunohistochemistry which showed robust RFP expression in tumor vasculature of *Atg12* ECKO tumors, while control tumors lacked RFP staining as expected (Supp. Fig. 5). To further test for phenotypes in *Atg12* ECKO animals in relation to the angiogenic switch, we also performed late deletion of *Atg12* at 10 weeks, which also did not alter *Atg12* ECKO survival relative to controls (p ≤ 0.7005, log rank test) (Fig. 7b). Tumor volume at 14 weeks was also not different between groups (ANOVA with Tukey’s test) (Fig. 7c). As expected, there was extensive staining present in *Atg12* ECKO animals but not control tumors. We also tested whether gross physiological features of insulin signaling were altered in *Atg12* ECKO mice, given the expansion of insulin-producing beta cells in *RT2-PNET* mice and increased insulin secretion. Comparing blood glucose levels at 14 weeks, we observed no statistical difference in non-starved blood glucose levels in *Atg12* ECKO versus controls (ANOVA/ Tukey’s test) (Supplemental Fig. 6). Similarly, body weight measurements in both male and female *RT2-PNET* mice showed no significant differences between groups (ANOVA/ Tukey’s test) (Supplemental Fig. 6).

Lastly, to test whether metastatic propensity was altered in this model, we enumerated the number of SV40+ metastatic tumor cells present in the livers of mice at endpoint (Fig. 6d). Interestingly, we uncovered a significant increase in SV40+ micro-metastasis in *Atg12* ECKO mice versus controls (p ≤ 0.0385, ANOVA) (Fig. 7e). Representative H&E of *RT2-PNETs* show grossly similar vascular patterns and tumor cell density between *Atg12* ECKO mice and controls (Fig. 7f, g). Thus, despite the lack of discernable differences in primary PNET progression or vascularity between the two cohorts, the loss of EC autophagy in RT2-PNETs results in increased micro-metastatic seeding in the livers of tumor-bearing mice.

## Discussion

Here, we dissect how the conditional genetic ablation of endothelial cell autophagy impacts *PyMT* breast tumor and *RT2-PNET* pancreatic neuroendocrine tumor progression. We show that VE-cadherin promoter Cre dependent ablation of *Atg12*, a gene required for autophagosome elongation, slows primary tumor growth in both orthotopic *PyMT* transplantation and autochthonous *PyMT* tumor models. A similar delay is observed with Atg5 ECKO, corroborating a autophagy-dependent phenotype[49]. This slower tumor growth phenotype is consistent with slowed growth observed in subcutaneously injected B16F10 melanoma cells into *Atg5* ECKO mice[25]. Most remarkably, we demonstrate increased metastasis in these mammary cancer models, which to our knowledge, has not been observed in previous studies of EC autophagy in tumor progression[25]. In addition, upon resection of primary PyMT tumors, we uncover increased rates of both primary tumor and metastatic recurrence. In contrast to the *PyMT* model, *Atg12* ECKO does not alter *RT2-PNET* primary tumor growth, but nonetheless, it does lead to an increase in liver micrometastasis, reinforcing an important functional role for EC autophagy in suppressing metastatic seeding and progression at distant sites in a second autochthonous tumor model.

Interestingly, these results resemble our recent studies of tumor cell autophagy in mammary cancer models, which demonstrate stark differences in the functional outcomes of autophagy inhibition in primary versus metastatic tumor growth[34, 50]. Overall, they highlight the exquisitely stage-specific and cancer-specific role of EC autophagy during tumor progression and metastasis. Nonetheless, our results do contrast with prior studies of melanoma in *Atg5* ECKO mice, which showed no differences in melanoma lung metastasis[25]. Although this may reflect fundamental differences between melanoma versus breast cancers and PNET, one consideration is that our studies employ models with slowed primary tumor progression kinetics compared to the aggressive B16F10 melanoma transplant model. As originally demonstrated in our studies of tumor cell autophagy during metastasis[34], the use of these slower progression tumor models may afford the ability to illuminate previously unrecognized roles for autophagy in the metastatic cascade. Remarkably, upon excision of primary PyMT tumors in *Atg12* ECKO mice, we observe higher frequencies of primary tumor and metastatic recurrence, and a dramatic increase in the percent of mice with lung metastasis at fixed time periods following the initial resection. Lastly, the introduction of *PyMT* cells into the systemic circulation of *Atg12* ECKO mice results in increased metastasis size, and a higher percentage of macro-metastasis in the lung. Altogether, our results support a clear functional role for EC autophagy in suppressing the outgrowth of tumor cells that exhibit an increased propensity for metastasis.

We examined primary *PyMT* tumors to identify tumor changes that may contribute to either slowed tumor growth or enhanced metastasis upon endothelial autophagy loss. Remarkably, there were no differences in the time to initially detect palpable tumors in either the orthotopic or autochthonous models; as a result, we hypothesize that *PyMT* tumors may be more sensitive to loss of endothelial autophagy in the mid-to-late stages of tumor progression. End stage *PyMT* tumors are highly heterogeneous, with necrotic regions of various sizes, tumor cell regions, and clusters of fibroblasts and stroma. While tumor cell proliferation, apoptosis, and necrotic areas were unchanged, tumors in *Atg12* ECKO mice did display increased levels of nuclear HIF1α, a marker of hypoxia-driven signaling and potential contributor to the metastatic phenotypes we have observed in response to the loss of EC autophagy [51]. As HIF1α has been linked to pro-metastatic lineage transitions, we scrutinized whether the increased in metastasis correlated with enhanced pro-metastatic basal differentiation that drives breast cancer metastasis in the setting of autophagy deficiency in tumor cells[52]. However, in contrast to genetic autophagy ablation in tumor cells, primary tumors in *Atg12* ECKO host animals exhibited no changes in either CK14 or CK5 expression[53], indicating that the expansion of basal sub-populations within the *PyMT* tumor compartment does not contribute to the increase in metastasis observed upon deletion of EC autophagy[46]. Delineating the role of HIF1α and other stress adaptation pathways in response to EC autophagy inhibition remains an important topic for future study.

We also scrutinized tumor growth and progression in the *RT2-PNET* pancreatic neuroendocrine model, which exhibit a temporally-defined angiogenic switch[54] and more robust responses to anti-angiogenic therapies[55] compared to breast cancer, where anti-angiogenic therapies have been less successful than anticipated[56]. In contrast to *PyMT*, *Atg12* ECKO mice did not exhibit changes in overall survival or in primary tumor volume at 14 weeks. Although the vasculature has a critical established role in *RT2-PNET* tumor progression, end stage *PyMT* tumors are much larger than *RT2-PNET* and exhibit significant necrosis, suggestive of unmet nutrient demands. Hence, we speculate that end stage *RT2-PNET* tumors may have a reduced requirement for EC autophagy. Nonetheless, liver micro-metastasis of *RT2-PNET* tumor cells is significantly increased in *Atg12* ECKO animals, consistent with the increased metastasis we observed in the *PyMT* models.

Remarkably, these results offer mechanistic insight into ongoing clinical trials addressing whether autophagy inhibition may be beneficial in breast and other cancers[2]. Our study indicates that although inhibiting EC autophagy may attenuate primary tumor growth, these benefits may be specific to certain cancer types, since primary progression was unaltered in the *RT2-PNET* model upon loss of EC autophagy. Additionally, our data broach that autophagy inhibition in endothelial cells may promote a HIF1α response in tumor cells that facilitates their adaptation to microenvironmental stresses and enables aggressive pro-metastatic behavior. Finally, our work suggests the importance of evaluating metastasis and optimizing therapeutic regimens in clinical trials to minimize the potentially detrimental effects of autophagy inhibition. Alternatively, the autophagy-independent functions of hydroxychloroquine on vessel normalization[25, 26] may afford significant therapeutic benefits and improve patient outcomes[57]. Rigorously interrogating whether the concomitant inhibition of angiogenesis and EC autophagy impacts cancer growth and metastasis remains an important question for future study.

## Author Contributions

This project was conceived and initiated in the lab of J.D. and continued in the lab of T.M. N.L.R. participated in experimental design, execution, analysis and manuscript writing and figure preparation; B.C. participated in experimental design, performance, statistical analysis, and figure preparation; S.B.E. assisted with histology; A.Q. assisted with experiments and analysis; A.G. and M.N. performed experiments; J.D. led project conception and generation of the mouse models, experimental guidance, manuscript revision and provided funding support; and T.M. led project conception, mouse studies, statistical analysis, manuscript writing and figure preparation, and provided funding support. J.D. Disclosures: None.

## Other Acknowledgements

We acknowledge support from the UCSF Histology and Biomarker Core Facility and the Flow Cytometry Core Facility at UCSF Parnassus. We acknowledge Bianca Charlene Rollbusch and Savannah Azzi, SDSU undergraduates, for support with quantification and imaging.

## List of abbreviations

Atg5: autophagy related protein 5
Atg12: autophagy related protein 12
CC3: cleaved caspase 3
CK5: cytokeratin 5
CK14: cytokeratin 14
EC: endothelial cell
ECKO: endothelial cell knockout
HUVEC: human umbilical vein endothelial cells
HIF1a: hypoxia inducible factor alpha
KI67: antigen Kiel 67
mTOR: mammalian target of rapamycin
PNET: pancreatic neuroendocrine tumor
PHH3: phosphor histone H3
PyMT: Polyoma Middle T
RT2-PNET: Rat insulin promoter T antigen 2 Pancreatic Neuroendocrine Tumor
RFP: red fluorescent protein
SV40: simian vacuolating virus 4

---

**Supplemental Fig. 1: FACS gating strategy testing for Cre-dependent RFP reporter.** Gating strategy shown for (**A**) non *Atg12* ECKO sample and (**B**) *Atg12* ECKO sample. To summarize, debris was gated out by examining forward scatter area versus side scatter area; singlets were gated in, live cells negative for Sytox Blue were gated in, then CD31-FITC+ gates established, and % RFP positivity of CD31% cells examined.

**Supplemental Fig. 2**: **Atg5 ECKO mice also display high Cre-dependent RFP reporter expression.** Immunohistochemistry showing increased P62 intensity of endothelial cells of *Atg5* ECKO mice in the mammary fat pad relative to *Atg5* fl/fl, tamoxifen controls. Scale bar = 50uM. **** indicates p≤0.0001.

**Supplemental Fig. 3: *PyMT* tumors lacking tumor cell Atg12 do not display altered HIF1A.** Left panel: representative image of end stage *Atg12 fl/fl* tumor; right panel: representative image of end stage Atg12 knockout tumor (*CagCreER; Atg12 fl/fl*).

**Supplemental Fig. 4: Autochthonous *Atg12* ECKO does not alter CK 5 expression in end stage *PyMT* tumors.** Quantification of mean fluorescent intensity (MFI) of CK5^+^ autochthonous end stage tumors showing no significant difference in *Atg12* ECKO tumors versus controls. One data point represents the average MFI per 10 high power fields per mouse (n=6 per genotype). Data are shown as mean ± SEM. Scale bar = 100uM.

**Supplemental Fig. 5: *Atg12* ECKO *RT2-PNET* tumors display Cre-dependent RFP expression at 14 weeks.** Immunohistochemistry for RFP Cre reporter of a representative control and *Atg12* ECKO *RT2-PNET* tumor at 14 weeks of age. Mice were administered tamoxifen or vehicle as indicated at 5 weeks of age (early deletion) to conditionally ablate autophagy or serve as controls. Scale bar= 5

**Supplemental Fig. 6: Body weight and blood glucose levels are similar in *Atg12* ECKO *RT2-PNET* mice.** Body weights of *Atg12* ECKO *RT2-PNET* females (**A**) and males (**B**) are similar to Cre- and vehicle treated controls at 14 weeks of age. Mice were administered tamoxifen or vehicle as indicated at ∼5 weeks of age (early). (**C**) Unfasted blood glucose levels, at 14 weeks at ∼2:00 pm were unaltered in *Atg12* ECKO *RT2-PNET* mice (males and females).

**Supplemental Table 1:** Immunofluorescence and immunohistochemistry primary antibodies used with company, catalog number, and dilution.

## References

[1] D. Hanahan, “Hallmarks of Cancer: New Dimensions,” Cancer Discovery, vol. 12, no. 1, pp. 31–46, 2022, doi: 10.1158/2159-8290.CD-21-1059.

[2] J. Debnath, N. Gammoh, and K. M. Ryan, “Autophagy and autophagy-related pathways in cancer,” Nature Reviews Molecular Cell Biology, vol. 24, no. 8, pp. 560–575, 2023/08/01 2023, doi: 10.1038/s41580-023-00585-z.

[3] F. Ferro, S. Servais, P. Besson, S. Roger, J.-F. Dumas, and L. Brisson, “Autophagy and mitophagy in cancer metabolic remodelling,” Seminars in Cell & Developmental Biology, vol. 98, pp. 129–138, 2020/02/01/ 2020, doi: 10.1016/j.semcdb.2019.05.029.

[4] Y. Wu et al., “Autophagy modulation in cancer therapy: Challenges coexist with opportunities,” European Journal of Medicinal Chemistry, vol. 276, p. 116688, 2024/10/05/ 2024, doi: 10.1016/j.ejmech.2024.116688.

[5] A. A.-O. DeMichele et al., “Targeting dormant tumor cells to prevent recurrent breast cancer: a randomized phase 2 trial,” (in eng), Nat Med., vol. 31, no. 1546-170X (Electronic), 10, pp. 3464–3474, 2025, doi: 10.1038/s41591-025-03877-3.

[6] Y. Chen and S. B. Gibson, “Three dimensions of autophagy in regulating tumor growth: cell survival/death, cell proliferation, and tumor dormancy,” Biochimica et Biophysica Acta (BBA)-Molecular Basis of Disease, vol. 1867, no. 12, p. 166265, 2021/12/01/ 2021, doi: 10.1016/j.bbadis.2021.166265.

[7] J. New et al., “Secretory Autophagy in Cancer-Associated Fibroblasts Promotes Head and Neck Cancer Progression and Offers a Novel Therapeutic Target,” (in eng), Cancer Res., vol. 77, no. 12, pp. 6679–6691, 12/1/2017 2017, doi: doi: 10.1158/0008-5472.CAN-17-1077.

[8] J. A. Rudnick et al., “Autophagy in stromal fibroblasts promotes tumor desmoplasia and mammary tumorigenesis,” Genes & Development, vol. 35, no. 13-14, pp. 963–975, 2021, doi: 10.1101/gad.345629.120.

[9] C. M. Sousa et al., “Pancreatic stellate cells support tumour metabolism through autophagic alanine secretion,” Nature, vol. 536, no. 7617, pp. 479–483, 2016, doi: 10.1038/nature19084.

[10] N. S. Katheder et al., “Microenvironmental autophagy promotes tumour growth,” Nature, vol. 541, no. 7637, pp. 417–420, 2017/01/01 2017, doi: 10.1038/nature20815.

[11] M. R. Spalinger et al., “PTPN22 regulates NLRP3-mediated IL1B secretion in an autophagy-dependent manner,” Autophagy, vol. 13, no. 9, pp. 1590–1601, 2017/09/02 2017, doi: 10.1080/15548627.2017.1341453.

[12] A. M. Leidal et al., “The LC3-conjugation machinery specifies the loading of RNA-binding proteins into extracellular vesicles,” Nature Cell Biology, vol. 22, no. 2, pp. 187–199, 2020, doi: 10.1038/s41556-019-0450-y.

[13] T. Monkkonen and J. Debnath, “Inflammatory signaling cascades and autophagy in cancer,” Autophagy, vol. 14, no. 2, pp. 190–198, 2018–02–01 2018, doi: 10.1080/15548627.2017.1345412.

[14] F. Jiang, “Autophagy in vascular endothelial cells.,” (in English), *Clinical and experimental pharmacology & physiology*, Review vol. 43,11, pp. 1021–1028, November, 2016 2016, doi: doi:10.1111/1440-1681.12649.

[15] B. A. Marzoog, “Endothelial Cell Aging and Autophagy Dysregulation,” (in English), *Cardiovascular & hematological agents in medicinal chemistry*, Review vol. 22,4, pp. 413–420, 2024, doi: 10.2174/0118715257275690231129101408.

[16] K. Torisu et al., “Intact endothelial autophagy is required to maintain vascular lipid homeostasis,” Aging Cell, vol. 15, no. 1, pp. 187–191, 2016/02/01 2016, doi: 10.1111/acel.12423.

[17] A.-C. Vion et al., “Autophagy is required for endothelial cell alignment and atheroprotection under physiological blood flow,” Proceedings of the National Academy of Sciences, vol. 114, no. 41, pp. E8675–E8684, 2017/10/10 2017, doi: 10.1073/pnas.1702223114.

[18] L. P. Bharath et al., “Impairment of autophagy in endothelial cells prevents shear-stress-induced increases in nitric oxide bioavailability,” Canadian Journal of Physiology and Pharmacology, vol. 92, no. 7, pp. 605–612, 2014/07/01 2014, doi: 10.1139/cjpp-2014-0017.

[19] T. Torisu et al., “Autophagy regulates endothelial cell processing, maturation and secretion of von Willebrand factor,” Nature Medicine, vol. 19, no. 10, pp. 1281–1287, 2013, doi: 10.1038/nm.3288.

[20] M. B. Schaaf, D. Houbaert, O. Meçe, and P. Agostinis, “Autophagy in endothelial cells and tumor angiogenesis,” Cell Death & Differentiation, vol. 26, no. 4, pp. 665–679, 2019/04/01 2019, doi: 10.1038/s41418-019-0287-8.

[21] N. Reglero-Real et al., “Autophagy modulates endothelial junctions to restrain neutrophil diapedesis during inflammation,” Immunity, vol. 54, no. 9, pp. 1989–2004.e9, 2021, doi: 10.1016/j.immuni.2021.07.012.

[22] H.-J. Wang, D. Zhang, Y.-Z. Tan, and T. Li, “Autophagy in endothelial progenitor cells is cytoprotective in hypoxic conditions,” American Journal of Physiology-Cell Physiology, vol. 304, no. 7, pp. C617–C626, 2013/04/01 2012, doi: 10.1152/ajpcell.00296.2012.

[23] D. Ge et al., “Identification of a novel MTOR activator and discovery of a competing endogenous RNA regulating autophagy in vascular endothelial cells,” Autophagy, vol. 10, no. 6, pp. 957–971, 2014/06/01 2014, doi: 10.4161/auto.28363.

[24] J. W. Xianjie Jiang, Xiangying Deng, Fang Xiong, Shanshan Zhang, Zhaojian Gong, Xiayu Li, Ke Cao, Hao Deng, Yi He, Qianjin Liao, Bo Xiang, Ming Zhou, Can Guo, Zhaoyang Zeng, Guiyuan Li, Xiaoling Li, Wei Xiong, “The role of microenvironment in tumor angiogenesis,” *Journal of experimental & clinical cancer research*, Review vol. 39,1, p. 204, September, 2020 2020, doi: 10.1186/s13046-020-01709-5.

[25] H. Maes et al., “Tumor Vessel Normalization by Chloroquine Independent of Autophagy,” Cancer Cell, vol. 26, no. 2, pp. 190–206, 2014, doi: 10.1016/j.ccr.2014.06.025.

[26] M. B. Schaaf et al., “Lysosomal Pathways and Autophagy Distinctively Control Endothelial Cell Behavior to Affect Tumor Vasculature,” (in eng), Frontiers Oncology, vol. 9, p. 171, March 20, 2019 2019.

[27] J. Verhoeven et al., “Tumor endothelial cell autophagy is a key vascular-immune checkpoint in melanoma,” EMBO Molecular Medicine, vol. 15, no. 12, p. e18028, 2023/12/07 2023, doi: 10.15252/emmm.202318028.

[28] H. A. Lane et al., “mTOR Inhibitor RAD001 (Everolimus) Has Antiangiogenic/Vascular Properties Distinct from a VEGFR Tyrosine Kinase Inhibitor,” Clinical Cancer Research, vol. 15, no. 5, pp. 1612–1622, 2009, doi: 10.1158/1078-0432.CCR-08-2057.

[29] P. M. v. Elizabeth Allen, Carmen M. Warren, Sadegh Saghafinia, Leanne Li, Mei-Wen Peng, Douglas Hanahan, “MetabolicSymbiosisEnablesAdaptiveResistanceto Anti-angiogenic TherapythatIsDependentonmTOR Signaling,” Cell Reports, vol. 15, no. 6, pp. 1144–1160, May 10, 2016 2016, doi: 10.1016/j.celrep.2016.04.029.

[30] A. Monvoisin, J. A. Alva, J. J. Hofmann, A. C. Zovein, T. F. Lane, and M. L. Iruela-Arispe, “VE-cadherin-CreER^T2^transgenic mouse: A model for inducible recombination in the endothelium,” Developmental Dynamics, vol. 235, no. 12, pp. 3413–3422, 2006, doi: 10.1002/dvdy.20982.

[31] T. Hara et al., “Suppression of basal autophagy in neural cells causes neurodegenerative disease in mice,” Nature, vol. 441, no. 7095, pp. 885–889, 2006/06/01 2006, doi: 10.1038/nature04724.

[32] R. Malhotra, E. Warne, Jp Fau-Salas, A. W. Salas, E Fau-Xu, J. Xu, Aw Fau-Debnath, and J. Debnath, “Loss of Atg12, but not Atg5, in pro-opiomelanocortin neurons exacerbates diet-induced obesity,“ (in eng), Autophagy, vol. 11, no. 1, pp. 145–54, 2015.

[33] H. Luche, O. Weber, T. NageswaraLRao, C. Blum, and H. J. Fehling, “Faithful activation of an extra-bright red fluorescent protein in “knock-in” Cre-reporter mice ideally suited for lineage tracing studies,” European Journal of Immunology, vol. 37, no. 1, pp. 43–53, 2007, doi: 10.1002/eji.200636745.

[34] T. Marsh et al., “Autophagic Degradation of NBR1 Restricts Metastatic Outgrowth during Mammary Tumor Progression,” Developmental Cell, vol. 52, no. 5, pp. 591–604.e6, 2020–03–01 2020, doi: 10.1016/j.devcel.2020.01.025.

[35] D. Hanahan, “Heritable formation of pancreatic β-cell tumours in transgenic mice expressing recombinant insulin/simian virus 40 oncogenes,” Nature, vol. 315, no. 6015, pp. 115–122, 1985/05/01 1985, doi: 10.1038/315115a0.

[36] C. T. Guy, R. D. Cardiff, and W. J. Muller, “Induction of Mammary Tumors by Expression of Polyomavirus Middle T Oncogene: A Transgenic Mouse Model for Metastatic Disease,” Molecular and Cellular Biology, vol. 12, no. 3, pp. 954–961, 1992, doi: 10.1128/mcb.12.3.954-961.1992.

[37] J. E. Fata et al., “The MAPKERK-1,2 pathway integrates distinct and antagonistic signals from TGFα and FGF7 in morphogenesis of mouse mammary epithelium,” Developmental Biology, vol. 306, no. 1, pp. 193–207, 2007/06/01/ 2007, doi: 10.1016/j.ydbio.2007.03.013.

[38] K. B. DeOme, L. J. Faulkin, Jr., H. A. Bern, and P. B. Blair, “Development of Mammary Tumors from Hyperplastic Alveolar Nodules Transplanted into Gland-free Mammary Fat Pads of Female C3H Mice*,” Cancer Research, vol. 19, no. 5, pp. 515–520, NP–NP, 1959.

[39] S. E. Martello, Xia, J., Kusunose, J., Hacker, B. C., Mayeaux, M. A., Lin, E. J., Hawkes, A., Singh, A., Caskey, C. F., & Rafat, M., “Ultrafast power doppler ultrasound enables longitudinal tracking of vascular changes that correlate with immune response after radiotherapy.,” Theranostics, vol. 14(18), pp. 6883–6896, 2024, doi: 10.7150/thno.97759.

[40] E. Y. Lin, Jones, J. G., Li, P., Zhu, L., Whitney, K. D., Muller, W. J., & Pollard, J. W., “Progression to malignancy in the polyoma middle T oncoprotein mouse breast cancer model provides a reliable model for human diseases,” *The American journal of pathology*, Scientific vol. 163(5), pp. 2113–2126, 2003 Nov 2003, doi: doi: 10.1016/S0002-9440(10)63568-7.

[41] E. Y. Lin et al., “Progression to Malignancy in the Polyoma Middle T Oncoprotein Mouse Breast Cancer Model Provides a Reliable Model for Human Diseases,” The American Journal of Pathology, vol. 163, no. 5, pp. 2113–2126, 2003, doi: 10.1016/s0002-9440(10)63568-7.

[42] A. Kuma et al., “The role of autophagy during the early neonatal starvation period,” Nature., vol. 432, no. 7020, pp. 1032–1036, 2004, doi: 10.1038/nature03029.

[43] C. C. Liao D, Seagroves TN, Johnson RS, “Hypoxia-inducible factor-1alpha is a key regulator of metastasis in a transgenic model of cancer initiation and progression.,” Cancer Res., vol. 67, 2, pp. 563–72, 2007.

[44] Michael S. Schieber and Navdeep S. Chandel, “ROS Links Glucose Metabolism to Breast Cancer Stem Cell and EMT Phenotype,” Cancer Cell, vol. 23, no. 3, pp. 265–267, 2013, doi: 10.1016/j.ccr.2013.02.021.

[45] A. Morandi, M. L. Taddei, P. Chiarugi, and E. Giannoni, “Targeting the Metabolic Reprogramming That Controls Epithelial-to-Mesenchymal Transition in Aggressive Tumors,“ Frontiers in Oncology, Review vol. Volume 7-2017, 2017. [Online]. Available: https://www.frontiersin.org/journals/oncology/articles/10.3389/fonc.2017.00040.

[46] G. Mondal, Gonzalez, H., Marsh, T., Leidal, A. M., Vlahakis, A., Phadatare, P. R., Bustamante Eguiguren, S., Bruck, M., Naik, A., Magbanua, M. J. M., Huppert, L. A., Wiita, A. P., Roose, J. P., Rosenbluth, J. M., & Debnath, J., “Autophagy-targeted NBR1-p62/SQSTM1 complexes promote breast cancer metastasis by sequestering ITCH,” Nature cell biology, vol. 27(7), pp. 1098–1113, 2025 Jul 2025, doi: 10.1038/s41556-025-01689-8.

[47] H. Maes et al., “Tumor Vessel Normalization by Chloroquine Independent of Autophagy,” Cancer Cell, vol. 26, no. 2, pp. 190–206, 2014, doi: 10.1016/j.ccr.2014.06.025.

[48] J. H. H. Masahiro Inoue, Napoleone Ferrara, Hans-Peter Gerber, Douglas Hanahan, “VEGF-A has a critical, nonredundant role in angiogenic switching and pancreatic beta cell carcinogenesis,” Cancer cell vol. 1, no. 2, pp. 193–202, March, 2002 2002, doi: 10.1016/s1535-6108(02)00031-4.

[49] F. Takeshita, K. Kobiyama, A. Miyawaki, N. Jounai, and K. Okuda, “The non-canonical role of Atg family members as suppressors of innate antiviral immune signaling,” Autophagy, vol. 4, no. 1, pp. 67–69, 2008/01/01 2008, doi: 10.4161/auto.5055.

[50] G. Mondal et al., “Autophagy-targeted NBR1–p62/SQSTM1 complexes promote breast cancer metastasis by sequestering ITCH,” Nature Cell Biology, vol. 27, no. 7, pp. 1098–1113, 2025/07/01 2025, doi: 10.1038/s41556-025-01689-8.

[51] G. L. Semenza, “The hypoxic tumor microenvironment: A driving force for breast cancer progression,” Biochimica et Biophysica Acta (BBA)-Molecular Cell Research, vol. 1863, no. 3, pp. 382–391, 2016/03/01/ 2016, doi: 10.1016/j.bbamcr.2015.05.036.

[52] K. J. Cheung et al., “Polyclonal breast cancer metastases arise from collective dissemination of keratin 14-expressing tumor cell clusters,” Proceedings of the National Academy of Sciences, vol. 113, no. 7, pp. E854–E863, 2016/02/16 2016, doi: 10.1073/pnas.1508541113.

[53] L. P. Schwab et al., “Hypoxia-inducible factor 1α promotes primary tumor growth and tumor-initiating cell activity in breast cancer,” Breast Cancer Research, vol. 14, no. 1, p. R6, 2012/01/07 2012, doi: 10.1186/bcr3087.

[54] M. Inoue, J. H. Hager, N. Ferrara, H.-P. Gerber, and D. Hanahan, “VEGF-A has a critical, nonredundant role in angiogenic switching and pancreatic &#x3b2; cell carcinogenesis,” Cancer Cell, vol. 1, no. 2, pp. 193–202, 2002, doi: 10.1016/S1535-6108(02)00031-4.

[55] R. Bill et al., “Nintedanib Is a Highly Effective Therapeutic for Neuroendocrine Carcinoma of the Pancreas (PNET) in the Rip1Tag2 Transgenic Mouse Model,” Clinical Cancer Research, vol. 21, no. 21, pp. 4856–4867, 2015, doi: 10.1158/1078-0432.CCR-14-3036.

[56] J. R. Mackey et al., “Controlling angiogenesis in breast cancer: A systematic review of anti-angiogenic trials,” Cancer Treatment Reviews, vol. 38, no. 6, pp. 673–688, 2012/10/01/ 2012, doi: 10.1016/j.ctrv.2011.12.002.

[57] S. Guelfi, K. Hodivala-Dilke, and G. Bergers, “Targeting the tumour vasculature: from vessel destruction to promotion,” Nature Reviews Cancer, vol. 24, no. 10, pp. 655–675, 2024/10/01 2024, doi: 10.1038/s41568-024-00736-0.

