## Supplementary figures and images for "Endothelial Cell Autophagy Suppresses Metastasis In Mouse Mammary and Pancreatic Neuroendocrine Tumor Models"

### Supplemental Figure 1

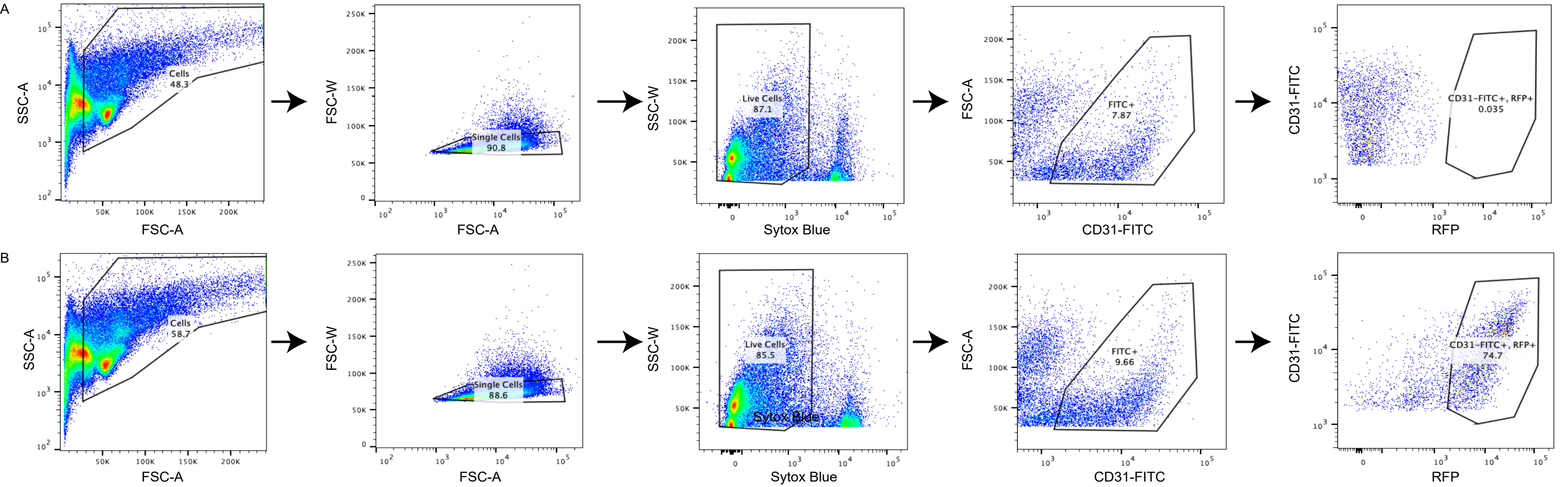

### Supplemental Figure 2

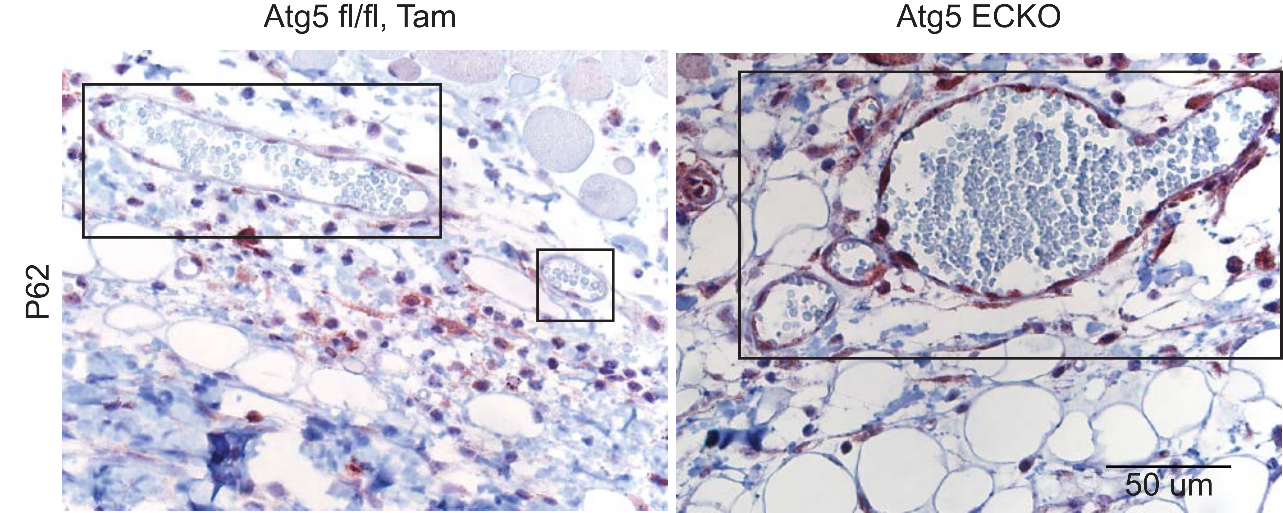

### Supplemental Figure 3

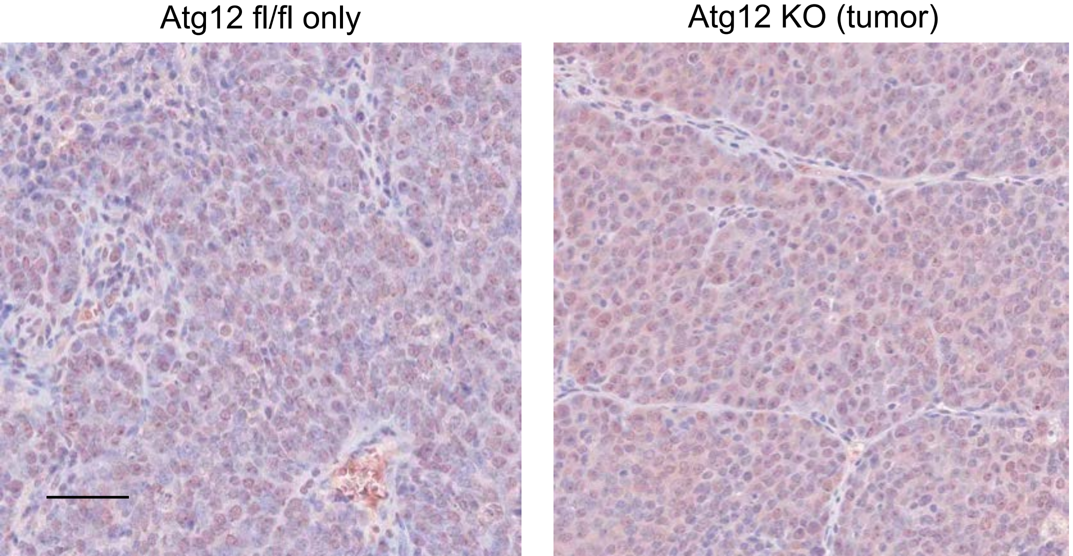

### Supplemental Figure 4

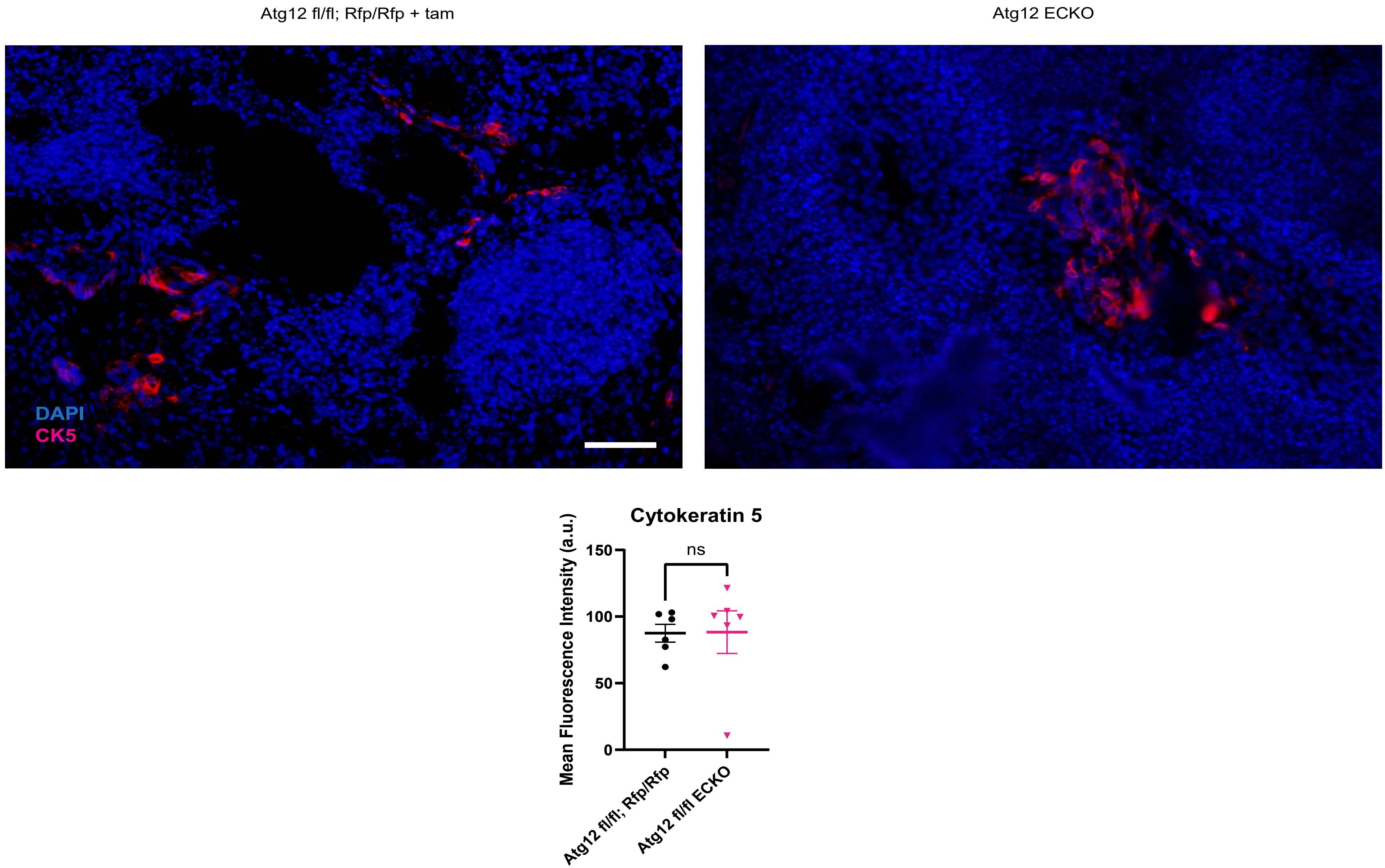

### Supplemental Figure 5

Representative Control

Atg12 fl/fl ECKO

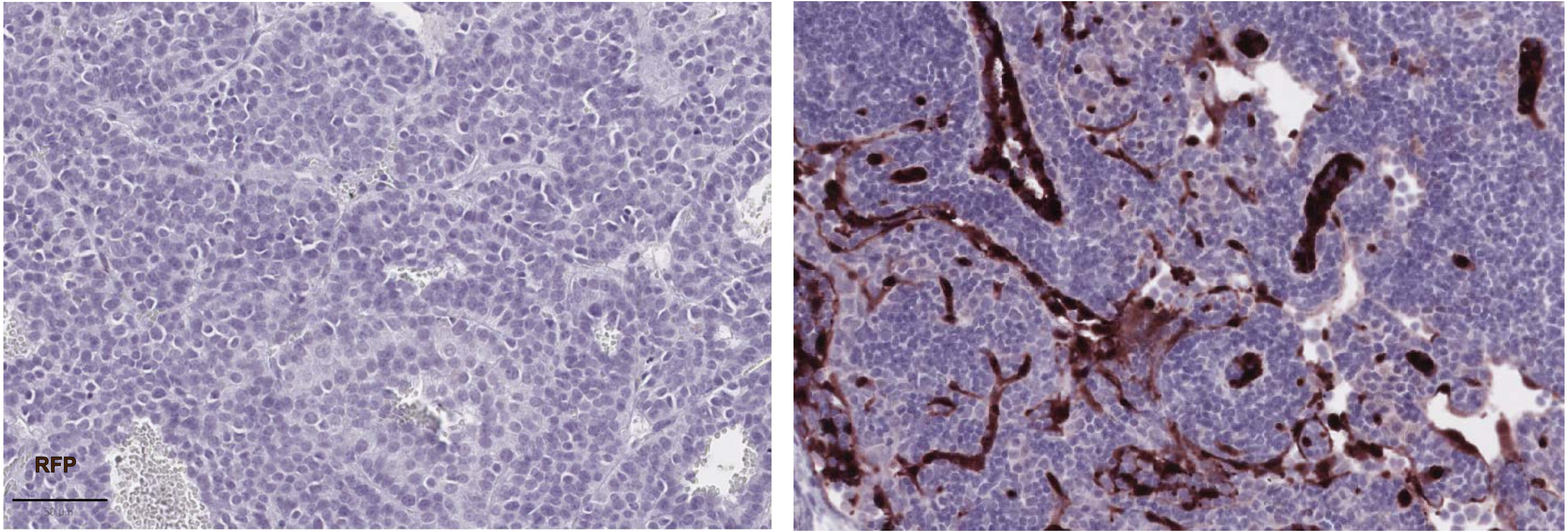

### Supplemental Figure 6

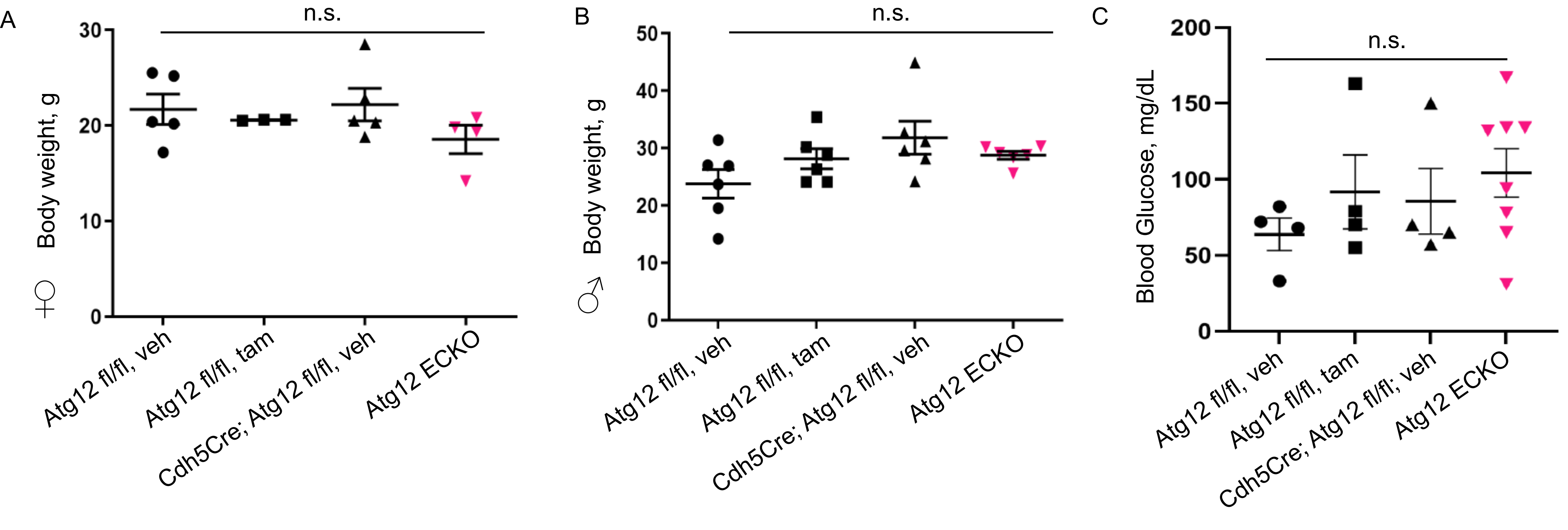
