## Supplemental Table 1 for "Endothelial Cell Autophagy Suppresses Metastasis In Mouse Mammary and Pancreatic Neuroendocrine Tumor Models"

| Target | Company (cat. no) | Concentration Used |
|---|---|---|
| Cleaved Caspase 3 | Cell Signaling (9661S) | 1:100 |
| Ki67 | Abcam (15580) | 1:100 |
| Phospho Histone H3 | Cell Signaling (9701) | 1:100 |
| CD31 (rabbit host) | Cell Signaling (77699) | 1:100 |
| CD31 (goat host, for CoIF) | R & D (AF3628) | 1:100 |
| CK14 | Abcam (ab7800) | 1:300 |
| SV40 | Santa Cruz (Sc147) | 1:50 |
| P62 | Progen (GP62-C) | 1:200 |
| CK5 | Cell Signaling (71536S) | 1:400 |
